# A novel quantitative trait locus implicates *Msh3* in the propensity for genome-wide short tandem repeat expansions in mice

**DOI:** 10.1101/2022.03.02.482700

**Authors:** Mikhail Maksimov, David G. Ashbrook, BXD Sequencing Consortium, Flavia Villani, Vincenza Colonna, Nima Mousavi, Nichole Ma, Abraham A. Palmer, Melissa Gymrek

## Abstract

Short tandem repeats (STRs) are a class of rapidly mutating genetic elements characterized by repeated units of 1 or more nucleotides. We leveraged whole genome sequencing data for 152 recombinant inbred (RI) strains from the BXD family derived from C57BL/6J and DBA/2J mice to study the effects of genetic background on genome-wide patterns of new mutations at STRs. We defined quantitative phenotypes describing the numbers and types of germline STR mutations in each strain and identified a locus on chromosome 13 associated with the propensity of STRs to expand. Several dozen genes lie in the QTL region, including *Msh3*, a known modifier of STR stability at pathogenic repeat expansions in mice and humans. Detailed analysis of the locus revealed a cluster of variants near the 5’ end of *Msh3*, including multiple protein-coding variants within the DNA mismatch recognition domain of MSH3, and a retrotransposon insertion overlapping an annotated exon. Additionally, gene expression analysis demonstrates co-localization of this QTL with expression QTLs for multiple nearby genes, including *Msh3*. Our results suggest a novel role for *Msh3* in regulating genome-wide patterns of germline STR mutations and demonstrate that inherited genetic variation can contribute to variability in accumulation of new mutations across individuals.

## Introduction

Studies of germline and somatic mutations have demonstrated considerable variation across individuals and species in both the rate and patterns by which mutations occur^1^. In some cases, this variation may be controlled by heritable factors influencing the function or expression of proteins involved in maintaining genome integrity. Indeed, genetic variants have been identified that disrupt DNA repair proteins^2,3^ and lead to “mutator” phenotypes in which affected individuals or cells accumulate specific types of mutations at a faster rate. While some of these phenotypes are highly deleterious, such as in cancer, common genetic variation can also result in more moderate mutator phenotypes that are only identified upon molecular interrogation^4^. Identifying genetic factors controlling this variation can give insight into mutation processes and DNA repair mechanisms.

Short tandem repeats (STRs), consisting of repeated sequence motifs of 1 or more bp, exhibit rapid mutation rates that are orders of magnitude greater than those for single nucleotide variants (SNVs)^5^. STR mutations typically result in expansions or contractions of one or more copies of the repeat unit. Expansion mutations are well known to cause a variety of disorders, including Huntginton’s Disease, hereditary ataxias, and myotonic dystrophy^6^. Further, we and others have recently implicated both small and large expansions and contractions at STRs in autism spectrum disorder^7,8^. Finally, somatic mutations at STRs, referred to as microsatellite instability (MSI), are a hallmark of certain cancer types^9^.

A large number of disease-focused studies have implicated proteins involved in mismatch repair (MMR) in regulating STR stability. For example, Lynch Syndrome, which results in predisposition to colon and other cancer types characterized by MSI, is caused by mutations that disrupt a variety of MMR proteins^10^. On the other hand, genome-wide association studies (GWAS) for age of onset of Huntington’s Disease have identified mutations in *MLH1*^*11*^ and *MSH3*^*12*^ that lead to increased somatic instability of the pathogenic trinucleotide expansion at *HTT*. Taken together, these studies suggest a critical role of MMR in regulating patterns of STR mutation.

The majority of studies of STR mutator phenotypes to date have focused on somatic repeat instability. However, studies of *de novo* STR and other mutation types have also demonstrated considerable variation in germline mutation rates across individuals^7,13^. While this variation is also potentially genetically controlled, this phenomenon is difficult to study in humans. Germline mutation rates are strongly confounded by parental age^14^, and mutation spectra may be influenced by environmental exposures^15^. Further, observed mutation patterns in children result from a mixture of mutation processes in the maternal and paternal germline. Thus, the relevant genetic variation controlling germline mutations could be harbored by either of the parents and is challenging to study in a typical GWAS setting.

Inbred mouse strains offer a unique opportunity to determine regulators of mutation processes because they can be used to study mutations that have accumulated over many generations under controlled settings. Further, within each strain offspring and both parents share essentially identical genomes, and thus we do not need to consider offspring and parental genotypes separately. We focused here on the BXD family^16^, which consists of more than 150 recombinant inbred (RI) strains that were generated by repeated inbreeding of the progeny of F_2_ crosses between inbred C57BL/6J and DBA/2J mice. We leveraged whole genome sequencing (WGS) for 152 BXD RI strains^17^ to perform quantitative trait loci (QTL) mapping for STR mutation phenotypes. This analysis revealed a novel QTL associated with the propensity of STRs to expand, and demonstrates how inherited genetic variation can affect the accumulation of new mutations across the genome.

## Results

### Identifying new mutations in BXD recombinant inbred mice

We previously built a reference set of autosomal 1,176,016 STRs with repeat units 2-20bp identified from the mm10 (C57BL/6J) reference assembly, and applied GangSTR^18^ to genotype these STRs using whole genome sequencing of 152 RI strains from the BXD cohort^17^. We used these genotypes to identify new STR mutations, which have presumably arisen over generations of breeding, by comparing the genotype at each RI strain to that expected based on the founder haplotype at that region. Heterozygous genotypes were removed since we determined these are likely enriched for STR genotyping errors. After filtering (**Methods**), we identified 18,135 STRs (1.5% of all STRs analyzed) for which at least one RI strain is homozygous for an STR length that does not match either of the founder genotypes, indicating a candidate new mutation (**Fig. 1a**). These mutations may occur at STRs that were previously fixed, or may occur at an STR that was already polymorphic in the founders. Tetranucleotide STRs represent the largest group of mutated loci, consistent with their abundance among the set of successfully genotyped loci (**Supplementary Fig. 1a**). Dinucleotide STRs, which are uniquely abundant in rodent genomes^19^, are under-represented in our dataset as a consequence of filtering due to low genotyping quality.

**Figure 1:**
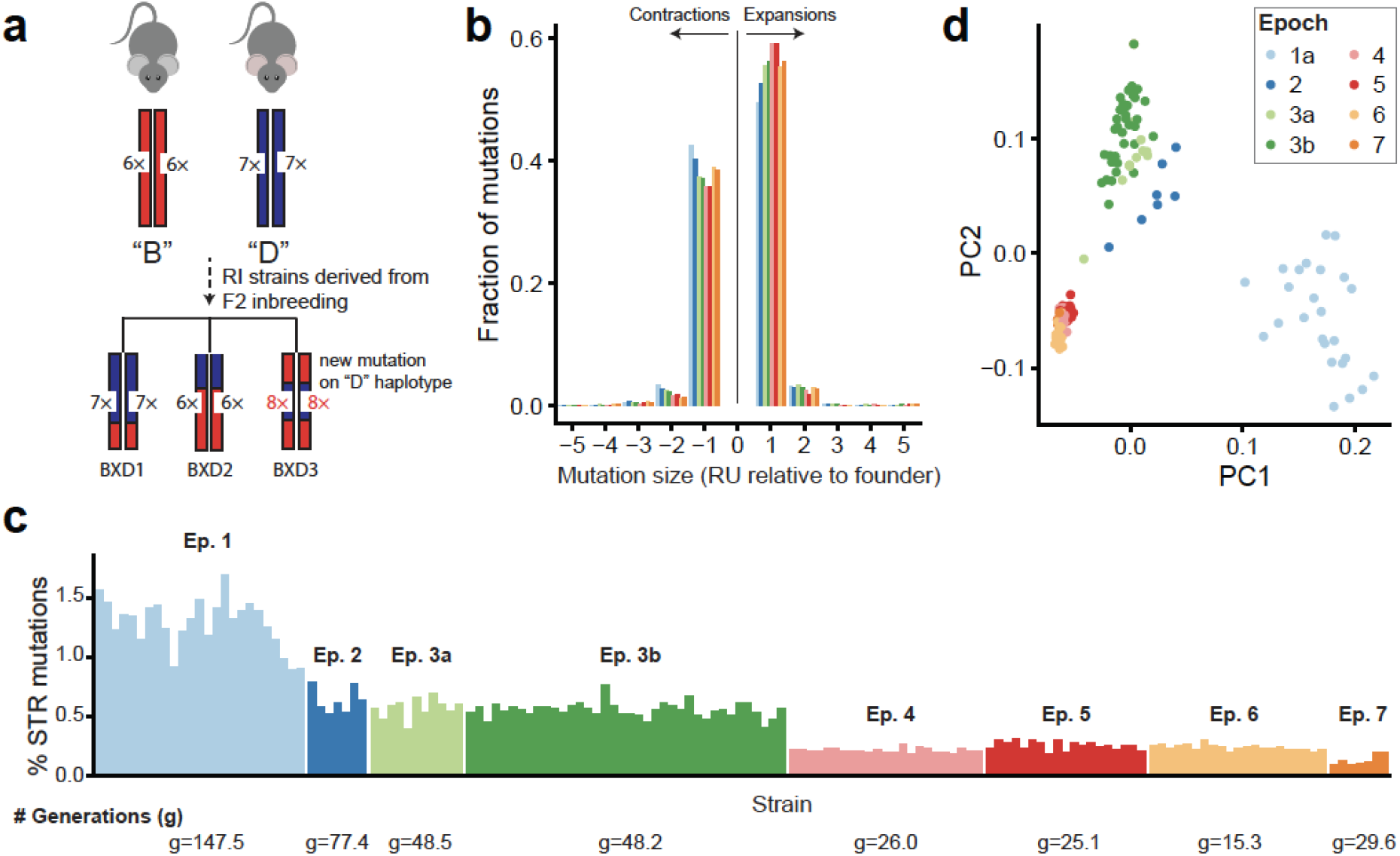
Characterizing new mutations in the BXD family. **a. Schematic of new mutation discovery in the BXD**. Each RI strain’s genome is a homozygous patchwork of segments derived from the founders, C57BL/6J (“B”, red) and DBA/2J (“D”, blue). Our STR mutation discovery pipeline considers a fixed set of STRs discovered in the mm10 reference panel (in the example shown, “B” has 6 copies and “D” has 7 copies of a particular STR). We identify new mutations as STRs with repeat lengths differing from the length of the founder inferred at that genome segment. In the example, strain BXD3 has a mutation to 8 copies that occurred on a haplotype inherited from the “D” founder. **b. Distribution of mutation sizes for each BXD epoch**. The x-axis shows mutation sizes in terms of the difference in number of repeat units from the founder allele. Positive sizes indicate expansions and negative sizes indicate contractions. Distributions are calculated separately for strains belonging to different epochs, indicated by bar color. **c. Percentage of genotyped STRs with a new variant for each RI strain**. New variants refer to any STR for which the observed allele does not match the expected founder allele. The average number of generations of breeding for strains in each epoch is annotated at the bottom of each panel. Strains are sorted by decreasing number of inbreeding generations within each epoch. **d. Principal component analysis (PCA) of new mutations**. PCA was performed on a binary matrix indicating whether each strain does or does not carry the new allele at each STR. The first two principal components separate strains by epoch indicating combinations of new mutations are shared among strains in each group. For **b-d**, colors denote BXD epochs.

We used SNP genotypes surrounding each STR to determine whether the mutation occurred on the haplotype originating from the “B” (C57BL/6J) vs. “D” (DBA/2J) founder, which enabled us to accurately determine the size of each mutation. We observed a slight excess of new mutations originating from the “B” founder (52.1%; **Supplementary Fig. 1b**), consistent with an overall slight excess of “B” haplotypes within BXD. The majority of mutations result in expansions or contractions of a single repeat unit compared to the founder with a bias toward expansion mutations (**Fig. 1b**). Mutations of two or more repeat units are slightly more prevalent among dinucleotide and trinucleotide repeats than among tetranucleotide repeats (**Supplementary Fig. 1c**). Both of these trends are consistent with those seen in human *de novo* STR mutations^7^. Nearly all mutations identified result in expansion or contraction by at most 5 repeat units, although our pipeline is not optimized to identify larger expansions.

Observed STR mutations are consistent with the known history of generation of the BXD strains. The BXD family is divided into epochs corresponding to various rounds of strain generation occurring over several decades and generated by multiple groups^16^. Assuming a constant mutation rate, the number of candidate STR mutations is expected to increase with the number of generations of breeding (**Fig. 1c**). While 54% of mutations identified are private to a single strain, the remainder are found in two or more strains (**Supplementary Fig. 2**). Principal components analysis based on genotypes at STRs for which we observe new mutations clearly separated strains by epoch (**Fig. 1d**), indicating many of these mutations are epoch-specific and likely arose in an individual ancestral to one or more epochs.

### Mapping quantitative trait loci for STR mutation phenotypes

We wondered whether observed differences in the number and size of mutations across strains could be driven by genetic variation affecting DNA repair or other pathways. To this end, we defined several quantitative phenotypes to summarize STR mutation patterns in each strain. We focused on three basic characteristics: *Mutation count* was computed as the fraction of genotyped STRs with a new mutation in each strain. Notably, this does not truly represent a germline *de novo* mutation rate, since observed mutations were homozygous and therefore must have occurred in ancestors to present-day individuals used for sequencing. *Mutation size* was calculated as the average change in repeat unit count, computed separately for expanded vs. contracted mutations in each strain. *Expansion propensity* was calculated as the fraction of new mutations in each strain for which the new allele is longer than the founder allele. For all phenotypes, we filtered new mutations seen in more than 10 strains, since those have likely been segregating within BXD on a variety of genetic backgrounds that differ from that of the individual where the mutation initially arose. These common mutations may also represent cases where the founder was incorrectly genotyped leading to false positive mutation calls. Due to their high mutation rates, recurrent mutations are expected, and so we did not restrict our analysis to mutations seen only once in our cohort. For mutation size and expansion propensity, we further restricted analysis to strains with at least 10 observed mutations since those phenotype values are unreliable when computed over a small number of mutations.

We performed genome-wide QTL mapping separately for each of these mutation phenotypes using R/qtl2^20^ and a set of 7,107 LD-pruned SNPs (**Fig. 2**). To account for population structure, R/qtl2 uses a linear mixed model with a kinship matrix generated using the leave-one-chromosome-out (LOCO) approach. The number of generations of inbreeding for each strain was used as a covariate. We determined genome-wide significance thresholds based on permutation analysis. QTL analysis did not identify any genome-wide significant loci for mutation size and found only a modestly significant signal on chromosome 7 for mutation count. However we identified a strong signal on chromosome 13 (max logarithm of the odds [LOD] = 8.7) associated with expansion propensity. Strains with the “B” haplotype at this locus tend to have higher expansion propensity than those with the “D” haplotype (**Fig. 2b**). This effect is consistent across most epochs (**Fig. 2c**), with the exception of later epochs for which a smaller number of mutations have accumulated. The QTL is centered around 90.4 Mb, but the 1.5-LOD support interval spans from 83.8-93.4 Mb, a region that encompasses several dozen genes (**Fig. 2d**).

**Figure 2:**
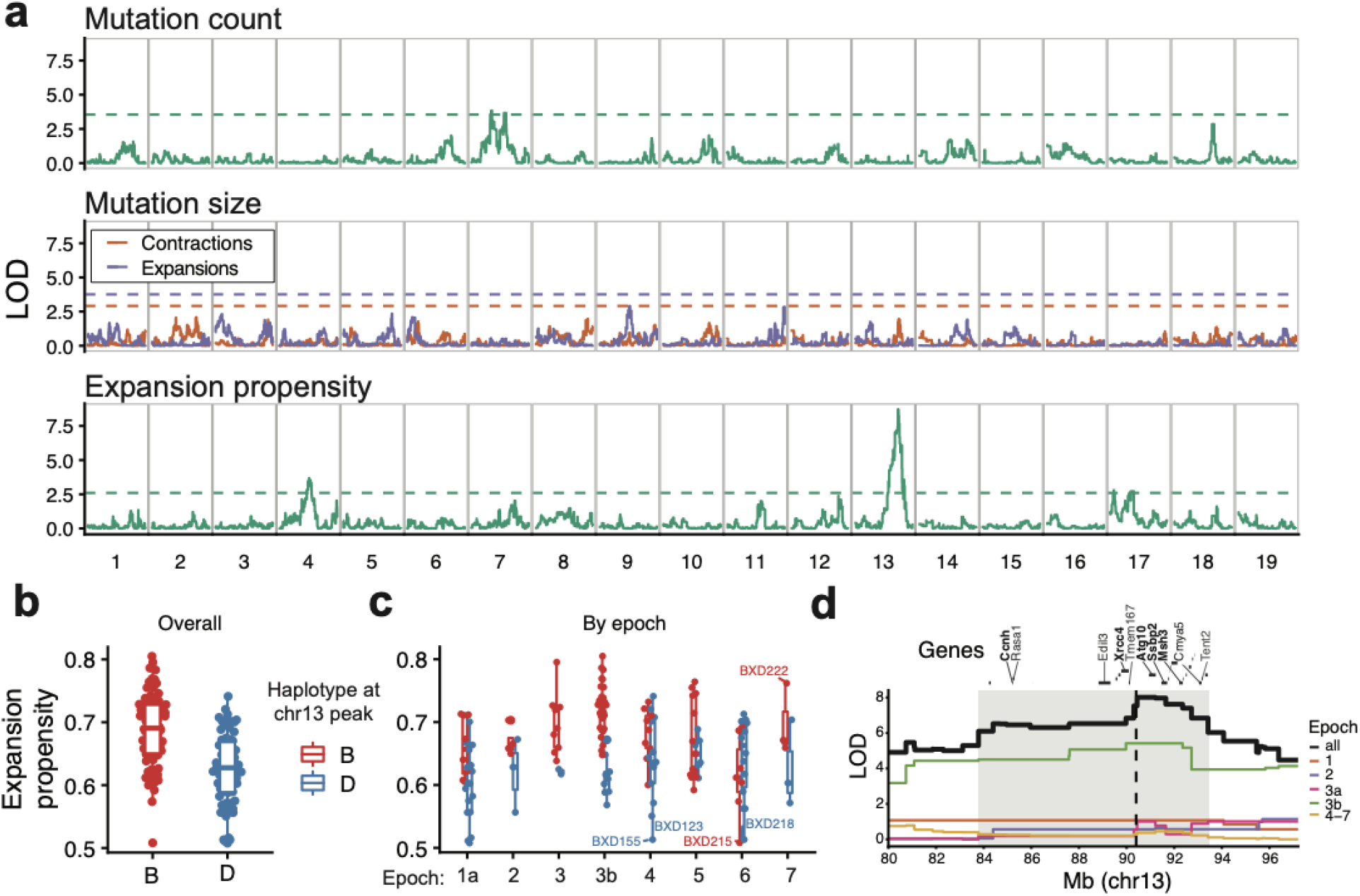
Discovery of QTLs for STR mutation phenotypes. **a. QTL mapping results** Panels show results for mutation count (top), mutation size (middle), and expansion propensity (bottom). The x-axis shows the genomic location, and the y-axis shows the LOD score of each SNP. For mutation size, red traces and blue traces represent contraction and expansion mutations, respectively. Dashed horizontal lines show genome-wide significance thresholds based on permutation analyses. **b-c. Increased expansion propensity is associated with the “B” haplotype at the chromosome 13 QTL**. Each point represents one strain. We used SNP haplotype blocks to assign each strain as harboring either the “B” (red) or “D” (blue) haplotype at this locus, shown separately for strains in each epoch. The y-axis denotes expansion propensity. Panel **b** shows the trend across all BXD strains, and panel **c** shows the trend separately for each epoch. Horizontal lines show median values, boxes span from the 25th percentile (Q1) to the 75th percentile (Q3). Whiskers extend to Q1-1.5*IQR (bottom) and Q3+1.5*IQR (top), where IQR gives the interquartile range (Q3-Q1). **d. Genes located in or near the QTL peak**. The y-axis shows the QTL signal (LOD score) for expansion propensity at chromosome 13 megabases 80-97. Black line=all strains, colored lines=QTL signal computed separately for strains in each BXD epoch. Horizontal bars denote a subset of genes near the center of the QTL peak. Genes known to be involved in DNA repair are highlighted.

To investigate whether the expansion propensity signal might be driven by specific types of STRs, we repeated QTL mapping separately for each repeat unit length. The signal is strongest by far for tetranucleotide STRs (**Supplementary Fig. 3**), which are the most abundant STR type in our dataset. Notably, all but tetranucleotide STRs have overall low mutation counts, resulting in unreliable estimates of expansion propensity for those categories (**Supplementary Fig. 4**). STRs with other repeat unit lengths show suggestive signals, but may be underpowered compared to tetranucleotides due to either low mutation counts or genotyping errors.

To test whether the chromosome 13 signal is influenced by our choice of filtering parameters, we repeated QTL mapping using a range of thresholds for the minimum number of mutations observed per strain and the maximum frequency of new mutations (**Supplementary Fig. 3**). Overall, the signal is robust to these filters and increases as we restrict analysis to successively rarer mutations. However, the signal is weaker when considering only private variants, which could be due to a combination of reduced power from lower mutation counts and enrichment of genotyping errors at private mutations.

Finally, we tested whether the observed signal replicates across BXD epochs, which were generated at separate times and locations, and could potentially have different environmental exposures or epoch-specific variants driving mutator phenotypes. The chromosome 13 signal is strongest in Epoch 3b (**Supplementary Fig. 5**), which has the most strains and therefore is the best-powered. However, other epochs show suggestive signals, and the signal is strongest when including all epochs. Thus, we concluded the causal variant(s) for this QTL are likely segregating within the entire BXD cohort and the QTL is unlikely to be due to an epoch-specific mutation or environmental phenomenon.

### Analysis of candidate variants disrupting protein-coding genes

We next sought to characterize the QTL on chromosome 13 for expansion propensity identified above. We first searched for variants in the region predicted to impact gene function. We identified 9,103 SNPs/indels and 160 STRs overlapping protein-coding genes. We additionally performed pangenome analysis to identify 3,698 large structural variants (SVs) (50bp<SV<10kbp; **Supplementary Fig. 6**). To reduce the search space, we removed rare variants (non-major allele fraction<0.15) and variants only weakly associated with the expansion propensity phenotype (model p-value <5e-4). We used the Ensembl Variant Effect Predictor (VEP)^21^ to annotate the predicted impact (modifier, low, moderate, or high) of the 8,213 variants that remained after filtering (**Supplementary Table 1; Supplementary Fig. 7**).

Based on previous studies of STR instability in cancer^10^ or modifiers of repeat expansion disorders^22^, we hypothesized that the observed STR mutator phenotype might be driven by variation in DNA repair genes. Of the genes in the QTL region, five are known to be involved in processes related to DNA repair: *Xrcc4* (non-homologous end joining to repair double strand breaks), *Ssbp2* (DNA damage response), *Ccnh* (nucleotide excision repair via the TFIIH complex^23^), *Atg10* (autophagy mediated effect^24^), and *Msh3* (involved in mismatch repair). Notably, *Msh3* has been widely implicated in STR stability. In humans, mutations disrupting *Msh3* function are known to contribute to MSI at tetranucleotide repeats in cancer^25^. On the other hand, *Msh3* is required for repeat expansions to occur^26,27^ and is a known modifier gene for STR stability in repeat expansion disorders^22^.

Of DNA repair genes in this region, *Ssbp2* and *Ccnh* contain only variants marked as modifiers by VEP which are unlikely to impact protein function directly, and *Xrcc4* contains multiple variants predicted to have low or moderate impact (**Supplementary Table 2**). *Atg10* has a more extensive variant profile with two moderate impact missense variants predicted as tolerated by SIFT^28^, one low impact synonymous variant and a multi-allelic coding sequence insertion (**Supplementary Table 2**) with a common allele resulting in an in-frame insertion (rs230013535) and a rarer allele causing a frameshift. Closer inspection of the frameshift allele revealed that all four strains carrying the allele are heterozygous and have lower genotype quality scores than other strains at the locus, suggesting this allele is a variant calling artifact and unlikely to explain the QTL signal.

*Msh3* contains the most variants with effects predicted by VEP, including one splice, four missense, and three synonymous mutations within protein-coding exons. Most of these are located within a variant-dense region in the 5’ end of the gene overlapping the mismatch recognition domain (**Supplementary Fig. 8, Supplementary Table 2, Fig. 3a**). One of the missense variants (rs48140189) is predicted by SIFT to be deleterious within a truncated transcript but is tolerated within both canonical transcripts. In addition to impactful variants within protein coding transcripts, we also identified three variants of interest mapping to a nonsense mediated decay (NMD) transcript of *Msh3* (ENSMUST00000190393). One of these is a structural variant corresponding to a 387bp insertion in C57BL/6J compared to DBA/2J (**Supplementary Fig. 6**). This type of insertion results from a long-terminal repeat (LTR) sequence left behind after an insertion of an IAPLTR2a retrotransposon into C57BL/6J and the subsequent deletion of its internal sequence following a recombination event^29^. The LTR spans nearly the entirety of exon 5 of an NMD transcript and falls between exons 4 and 5 of the canonical transcript (**Supplementary Figure 8b; Fig. 3b**). Additionally, a pair of SNPs (rs49933543; rs48930870) are predicted to reside within the 3’ splice acceptor site of this 4th intron of the NMD transcript and are well correlated with the mutator phenotype. We further examined other SVs within each gene that passed the association and allele frequency criteria regardless of their impact predicted by VEP (**Supplementary Table 3**). *W*hile *Atg10, Ssbp2* and *Xrcc4* harbor several large (>50bp) structural variants, neither of these is predicted to overlap with any meaningful feature. Moreover, with the exception of a 215bp deletion within *Atg10*, the *Msh3* LTR variant is most strongly associated with the mutator phenotype. Overall, given its known role in STR stability and the high density of variants with predicted impact overlapping its mismatch recognition domain, our results suggest *Msh3* as a strong candidate gene for this QTL.

**Figure 3:**
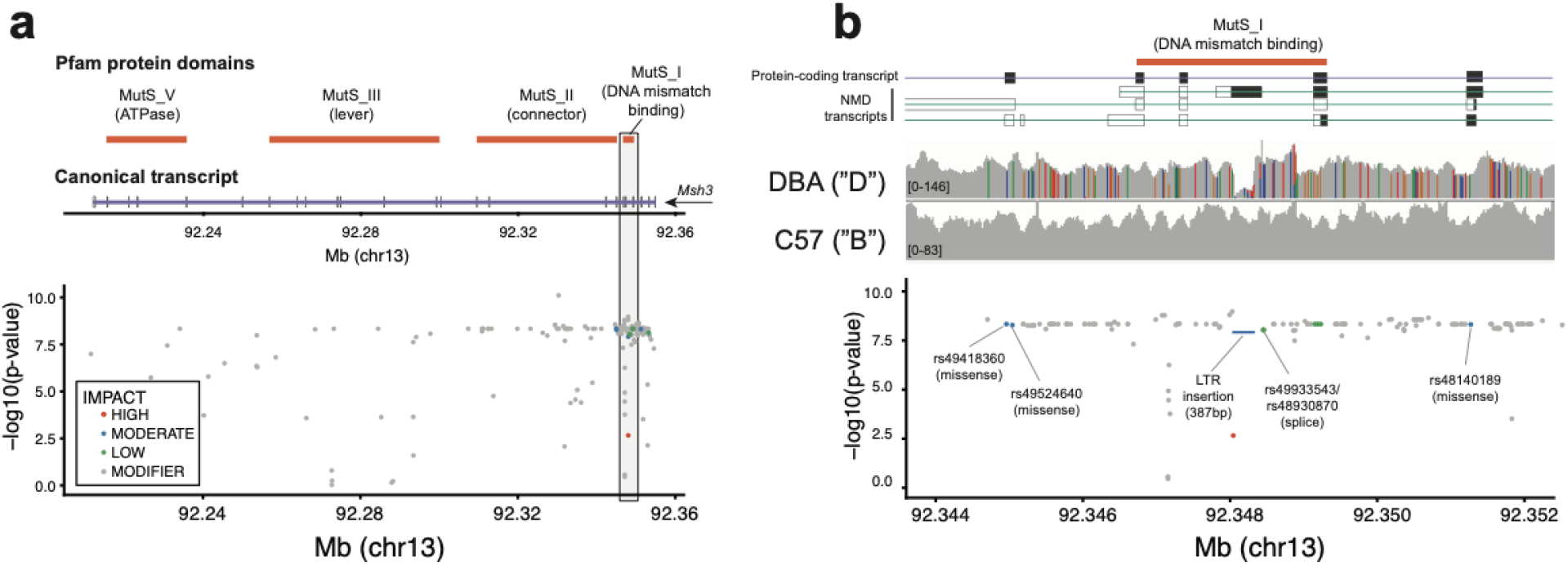
Variants predicted to impact *Msh3*. **a. Summary of variants overlapping *Msh3***. The top panel shows the canonical protein-coding transcript of *Msh3* (purple) and protein domains (orange rectangles) obtained from Pfam^30^. The bottom panel shows the location (mm10; x-axis) of variants and their association with the expansion propensity phenotype (-log10 p-values; y-axis). Variants are colored by their impact predicted by VEP (high=red; moderate=blue; low=green; modifier=gray). **b. Summary of variants in the variant-dense 5’ region of *Msh3***. Top and bottom panels are the same as in **a**. The middle panel shows a histogram of read coverage as visualized using the Integrative Genomics Viewer^31^. Gray bars denote positions that match the reference genome, which is based on C57BL/6J. Colored bars denote mismatches from the reference.

### Expansion propensity QTL co-localizes with multiple *cis*-eQTLs

We next wondered if the QTL for expansion propensity might also be mediated through *cis*-regulatory variants affecting expression of genes in this region. To this end, we compiled 54 publicly available gene expression microarray datasets encompassing 30 tissues (**Supplementary Table 4**) with sample sizes ranging from 11-79 strains. While the datasets were acquired using multiple microarray platforms, under different experimental conditions, and across a range of tissues, we find overall that *Ccnh* and *Ssbp2* are among the most highly expressed genes within the QTL region, *Msh3* has average expression, while *Atg10* and *Xrcc4* are expressed slightly below average (**Supplementary Fig. 9**). For downstream analyses, we restricted to 40 expression datasets with at least 30 strains (**Supplementary Fig. 10**).

For each remaining dataset, we performed a separate expression QTL (eQTL) analysis for 26 protein coding genes for which expression levels are available in at least half of the datasets (**Supplementary Fig. 11**). We considered only probes not overlapping SNPs for comparing gene expression levels and used the number of variants per probe as a covariate in eQTL mapping to avoid confounding the true variability with differences in probe hybridization efficiency. Notably, this excluded a large number of probes for *Msh3* since many overlap multiple SNPs in the highly variable 5’ end of the gene (**Supplementary Fig. 12**). We then ranked genes by the proportion of datasets where the maximum eQTL LOD exceeded the permutation-based threshold for significance (**Supplementary Fig. 13**). We observed robust eQTL signals for *Ssbp2, Ag10* and *Ccnh* in 29, 18 and 14 datasets respectively. We also found eQTL signals for *Xrcc4* and *Msh3*, albeit in a smaller number of datasets: 6 and 4 respectively (**Fig. 4a, Supplementary Fig. 14a**). The eQTL for *Atg10* shows the most consistent colocalization with the QTL peak across datasets (**Supplementary Fig. 14a)**. However, eQTLs for most genes in the region are strongly co-localized with the QTL (**Fig. 4a)**, making it difficult to prioritize a single causal gene based on the eQTL signal alone.

**Figure 4:**
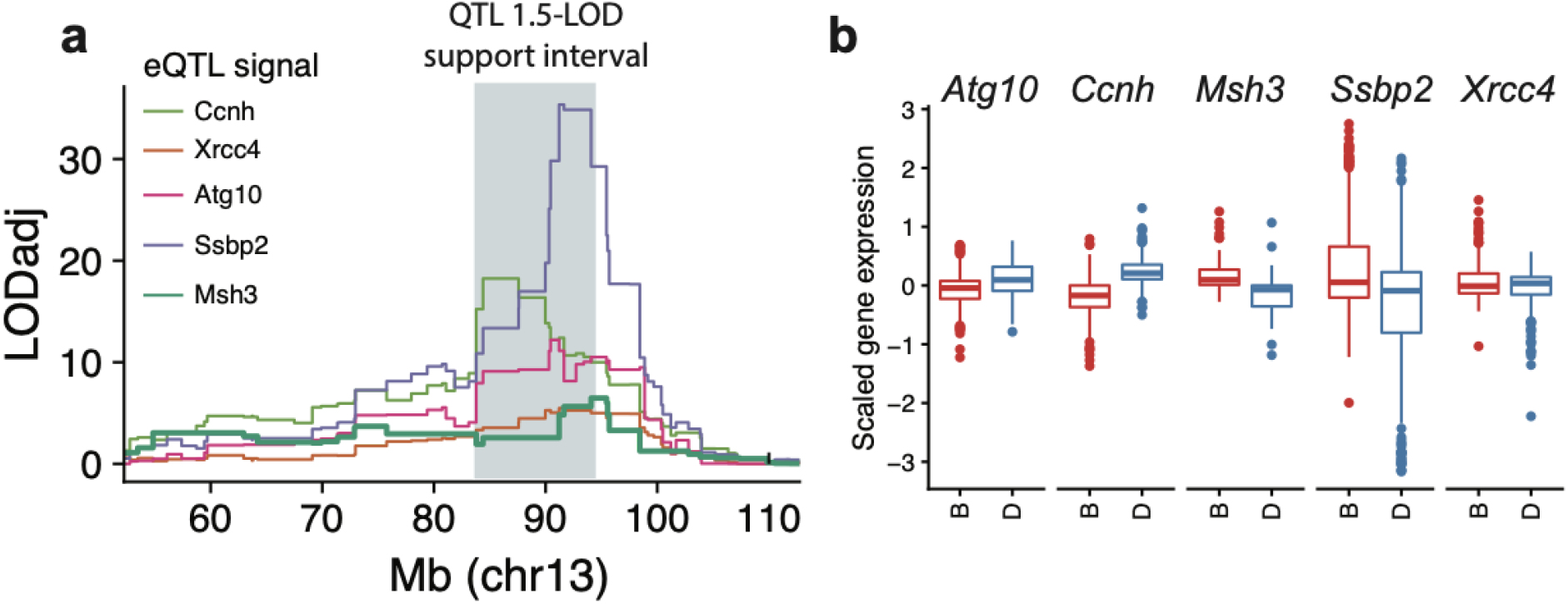
A QTL for expansion propensity on chromosome 13 colocalizes with eQTLs for multiple DNA repair genes. **a. Co-localization of expansion propensity and eQTL signals**. Colored traces denote eQTL LOD scores. Each line shows the expression dataset with the strongest eQTL for that gene. eQTL LOD scores were adjusted for multiple hypothesis testing for each gene based on the number of probes tested. The gray shaded box shows the 1.5-LOD support interval for the expansion propensity QTL. **b. Distribution of gene expression for strains with “B” vs. “D” haplotypes**. Panels show gene expression for each gene for strains assigned the “B” (red) vs. “D” (blue) haplotypes at the QTL locus. Data shown is aggregated across all GeneNetwork datasets with a significant eQTL for each gene. Distributions per dataset are shown in **Supplementary Fig. 14b**.

In all tissues with a significant eQTL for *Msh3*, we observed a consistent direction of effect, with higher *Msh3* expression for strains carrying the “B” haplotype associated with increased expansion propensity (**Fig. 4b, Supplementary Fig. 14b**). Detailed analysis of the *Msh3* eQTL shows that the signal is strongest when considering probes and variants in the 5’ end, even after adjusting for hybridization efficiency due to SNPs in this region (**Methods**; **Supplementary Fig. 15**). This result is consistent with previous studies in humans, in which increased *Msh3* expression driven by polymorphism in the 5’ end of the gene was associated with increased somatic instability at the trinucleotide repeat involved in Huntington’s disease^12^.

Finally, we examined tissue-specific expression of each of the candidate DNA repair genes using the Bgee^32^ database (**Supplementary Table 5**). *Msh3* is most highly expressed in oocytes, where de novo mutations are known to arise^33^. On the other hand, *Ssbp2* and *Atg10*, which are closest to the QTL center, are most highly expressed in brain and heart structures, respectively, which are unlikely to be relevant for germline mutations. *Ccnh* is expressed in a variety of tissues including female gonads, and *Xrcc4* is expressed in spermatocytes and oocytes. However, variants overlapping those genes have lower LOD scores for association with expansion propensity than variants overlapping *Msh3* or *Atg10* (**Fig. 2d, Supplementary Fig. 7**).

## Discussion

Genetic variation impacting proteins involved in DNA repair processes have the potential to drive genome-wide variation in mutation rates and patterns across individuals of a species, both in the context of disease but also across healthy individuals. Identifying these determinants may give insights into disease risk or progression and could improve population-genetic models of mutations. Recombinant inbred (RI) strains such as those in the BXD cohort have accumulated mutations over dozens of generations of inbreeding, offering a unique opportunity to map genetic determinants of these “mutator phenotypes”. Here, we performed QTL mapping for three quantitative STR mutator phenotypes, and identified a robust QTL on chromosome 13 for expansion propensity in mice. The QTL region encompasses 30 protein-coding genes, including *Msh3*, an important component of the DNA mismatch repair (MMR) machinery^3^.

*Msh3* is well-known to be involved in regulating STR stability. *Msh3* is one of six homologs of the *E. coli* MutS MMR protein, and heterodimerizes with *Msh2* to recognize and repair long insertion or deletion loops that arise during DNA replication^34^, often due to misalignment of strands at STR regions. Model organism studies have demonstrated that *Msh3* is required for the formation of pathogenic repeat expansions^26,27^. Further, inherited variants in *Msh3* have been reported to modify age of onset and severity of Huntington’s Disease^22^, and a polymorphism in the 5’ end of *Msh3* has been associated with increased *Msh3* expression and somatic instability of the trinucleotide repeat implicated in Huntington’s Disease^12^. Finally, mutations disrupting *Msh3* are often found in cancers exhibiting microsatellite instability^25^.

Here, we report a novel role for *Msh3* in altering genome-wide germline mutation patterns at STRs. Previous studies have focused either on *Msh3*’s role as a modifier gene for the stability of large repeat expansions at a small number of well-studied loci, or on its connection with somatic STR instability in cancer. Our results suggest that in addition to these roles in disease, common mutations affecting *Msh3* may contribute to biases in mutation sizes in the germline at the hundreds of thousands of short STRs across the genome. Similar to previous findings in inbred mice^26^, we find evidence that both protein-coding sequence variants overlapping the DNA mismatch recognition domain, as well as *Msh3* expression levels, could collectively contribute to the increased expansion propensity in mice harboring the “B” vs. “D” haplotypes at this locus (**Fig. 5**). Notably, we found that these haplotypes are shared with other inbred strains commonly used in mouse genetics (**Supplementary Fig. 16**), suggesting they could drive variation in STR mutation profiles in other mouse cohorts.

**Figure 5:**
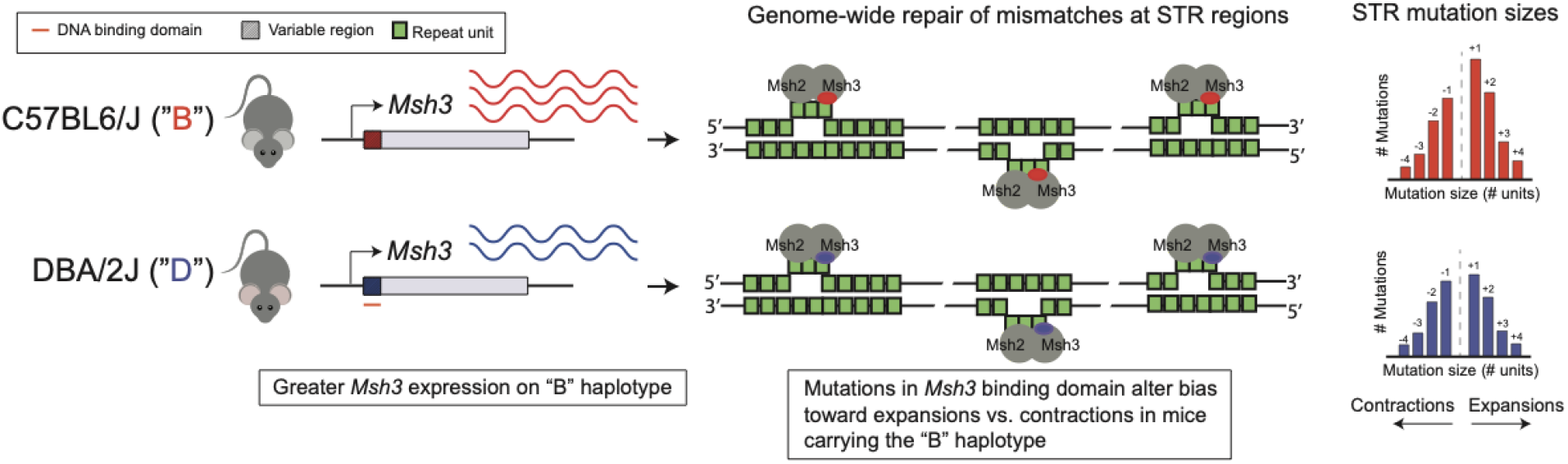
Schematic overview of proposed mechanisms for the expansion propensity QTL. BXD mice carrying the “B” haplotype at the chromosome 13 QTL locus tend to have higher *Msh3* expression than those carrying the “D” haplotype. *Msh3* and *Msh2* form the heterodimer MutSβ which recognizes strand misalignments, such as those formed by STRs (repeat units shown in green), across the genome during DNA replication. A cluster of sequence variants at the 5’ end of *Msh3* overlapping the DNA binding domain may alter how the heterodimer to recognizes and corrects misalignments at STRs, leading to a bias in expansion vs. contraction mutations in mice carrying the “B” haplotype at this locus.

Identifying a single causal gene or variant in the QTL locus identified is challenging in the BXD family, which harbors long unbroken haplotypes spanning several megabases^16^. The abundance of literature evidence regarding the role of *Msh3* in STR stability in other contexts, and the high density of variants in the key region of the protein important for recognizing mismatched DNA, strongly suggests it as a causal gene for this locus. However, we could not definitively rule out a role for other genes in this region. In particular, *Atg10* falls closest to the center of the QTL peak, and eQTL signals for *Atg10* are most consistently co-localized with the QTL. We additionally identified multiple protein-coding variants and an SV overlapping this gene. However, *Atg10* has only been indirectly connected with DNA repair through the autophagy system^35^. Further, whereas *Msh3* is most highly expressed in oocytes where germline mutations are likely to arise, *Atg10* is most highly expressed in heart and other structures less likely to be related to a mutator phenotype.

In addition to multiple protein-coding variants that have been previously reported^26^, our analyses revealed a 387bp indel, near the 5’ end of the gene and falling between exons 4 and 5 which encode the DNA mismatch recognition domain. This indel is due to an IAP LTR insertion on the “B” (C57BL6/J) which is missing in “D” (DBA/2J) and many other classic mouse strains (**Supplementary Fig. 16**), and forms an exon of a non-canonical transcript of *Msh3*. IAP LTRs are one of the few active retrotransposon families in the mouse genome^36^. Two of the most well-studied variants in mice have arisen through IAP LTR insertion: Agouti Viable Yellow^37^ and Axin fusion^38^. While IAP LTR elements are typically heavily methylated^39^, the element at this locus is a member of the IAPLTR2a group, which are overrepresented among hypomethylated LTRs^40^, harbor transcription factor binding sites which can potentially contribute to regulation of nearby genes^41^, and have been shown to induce alternative splicing of nearby exons^36^. Further, retrotransposons including IAP elements, have been shown to regulate transcription of their host genes in oocytes^42^, where Msh3 is most highly expressed.

Our study faced multiple limitations. First, our pipeline is not optimized to identify mutations resulting in large repeat expansions, such as those implicated in Huntington’s Disease. Additionally, available WGS data was generated using PCR+ protocols, and many STRs, in particular error-prone dinucleotide repeats, had to be filtered from our analysis due to low-quality calls. Second, while we analyzed gene expression data for dozens of tissues collected for BXD samples, we did not have access to expression data for tissues most relevant for germline mutation processes. In future studies, data from tissues such as oocytes or sperm might give further insights into gene regulatory mechanisms driving this signal. Finally, experimental validation of individual causal genes or variants for this phenotype is challenging: the STR mutation phenotypes measured here are based on mutations that have arisen over decades of inbreeding, and would not be evident in genome-edited cell lines or animals observed for a small number of generations.

In summary, our study reveals a novel QTL for STR mutation patterns, providing a striking example of the influence of inherited variants on germline mutation properties. Additional modifiers for both STR and other mutator phenotypes are likely to exist in humans or in other model organism datasets. We anticipate that further investigation of these mutation modifiers will provide new insights into mutation processes both in health and disease.

## Methods

### Whole genome sequencing of the BXD cohort

Genome-wide STR and SNP genotypes for 152 RI strains and the two BXD founders, C57BL/6J (“B”) and DBA/2J (“D”), were previously generated from whole genome sequencing data from the BXD Sequencing Consortium^17^. Epoch labels were obtained from Ashbrook, et al.^16^.

### SNP marker maps for founder inference and interval mapping

We prepared a marker-by-strain matrix of founder labels (“B” vs. “D”) for BXD RI strains using SNP genotypes at 7,124 autosomal LD-pruned markers published on GeneNetwork (**URLs**). For SNPs not directly genotyped from WGS in the BXD, we chose the next closest SNP based on genomic distance that was less than 500 Kbp away. In a small number of cases, the closest SNP was the same for multiple markers, in which case a single marker/snp combination was retained producing a final list of 7,107 markers. R/qtl2^20^ version 0.24 was used to calculate founder genotype probabilities suitable for QTL mapping using the ‘calc_genoprob’ function with default parameters. We then generated a complete list of SNP founder labels with maximum marginal probabilities using the ‘maxmarg’ function with ‘minprob’ parameter set to 0.5. Founder labels at individual markers were used to find start and stop positions of haplotype blocks using a connected components clustering approach (R tidygraph).

### Identifying and phasing new STR mutations

We identified candidate STR mutations as STR genotypes in RI strains not matching genotypes in either of the two founder strains. In cases where one or both founders were not directly genotyped, we first inferred missing STR calls in founders (below). We intersected each candidate new mutation with haplotype blocks inferred from SNPs to assign each mutation as occurring on the “B” vs. “D” haplotype. STRs falling in a gap between blocks were assigned to the nearest block. We excluded new variants where either the RI or founder strain were heterozygous, which likely indicates either poor quality STR genotypes or incomplete inbreeding at that locus. Finally, we excluded strain BXD194, in which we found an outlier number of new mutations (>2-fold higher than other strains in the same epoch), from downstream analyses.

### Inferring missing founder STR genotypes

We used R/qtl2 to infer missing founder STR genotypes from genotypes observed in RI strains. First, we imputed founder labels (“B” or “D”) for each STR genotype in the RI strains. For the subset of loci at which both founder strains were genotyped and did not share a common allele, we could unambiguously assign “B” or “D” genotype labels to each genotyped RI strain. RI strains with genotypes not matching either founder were assigned missing labels. For the remaining polymorphic loci missing at least one founder genotype, we could not directly infer the founder label and initially set all genotypes at those loci to missing values. We used the R/qtl2 ‘interp_map’ function to interpolate linkage distances between STRs from physical and genetic SNP marker maps at the 7,107 LD-independent markers described above. We then used R/qtl2 functions ‘calc_genoprob’ followed by ‘maxmarg’ to impute missing founder labels. Then, for each STR with a missing founder genotype, we determined the distribution of repeat lengths in strains inferred to have the corresponding founder label at that locus. If at most one RI strain had a genotype differing from the modal repeat length, the founder was inferred to have the modal allele. Otherwise, the locus was removed from downstream analysis.

### Characterization of new STR mutations

We performed principal components analysis (PCA) to characterize sharing of new mutations across strains. We constructed a strain-by-locus matrix of indicator values indicating the presence (1) or absence (0) of a new STR genotype in each strain at each locus. We then performed PCA using the builtin ‘prcomp’ function in R with centering but without scaling.

### Computing STR mutator phenotypes

We calculated three separate mutator phenotypes for each strain. *Mutation count* was calculated as the number of STRs with new mutations divided by the number of successfully genotyped loci in that strain. *Mutation size* was calculated as the average difference in repeat count between the new genotype and the founder genotype at each mutation. Mutation size was computed separately for expansion and contraction mutations. *Expansion propensity* was calculated as the fraction of new mutations in each strain for which the RI genotype was longer than the founder genotype. Unless otherwise noted, we removed STR mutations seen in 10 or more strains, as those likely do not represent new mutations.

### QTL mapping for STR mutator phenotypes

QTL mapping for each mutator phenotype was performed based on the set of LD-pruned SNPs described above using a linear mixed model approach implemented in R/qtl2. Each phenotype was analyzed separately. We used the “calc_kinship” function to prepare a strain relatedness matrix using the leave-one-chromosome-out (LOCO) method. In addition to supplying a vector of phenotype values, genotype probability and kinship matrices, we also input a vector of the number of inbreeding generations as a covariate. We used ‘scan1perm’ to calculate permutation-based genome-wide significance thresholds based on 100 permutations. For each QTL analysis performed, strains with fewer than 10 total new mutations were excluded from analysis since they produce noisy mutator phenotype values.

### Variant annotation

The initial set of variants for annotation analysis contained 166,681 SNPs and 2,355 STRs genotyped previously in the BXD cohort^17^ and located between the boundaries of the confidence interval for the QTL on chromosome 13. We additionally obtained structural variants (SVs) based on pangenome analysis (see below). After filtering for variants within protein coding genes in the QTL region based on the GENCODE M25 release gene annotations, 46,320 SNPs, 736 STRs and 7,623 SVs remained. SVs smaller than 50bp were removed leaving 983 SVs. After filtering for only segregating variants and removing variants where more than half the strains had a missing value, 9,103 SNPs, 160 STRs and 983 SVs remained. Non-major allele frequency was calculated for each variant as the proportion of alleles at the locus that were not the most abundant allele after removing strains with missing genotypes. We used VEP^21^ v103.1 with the Ensembl cache v102 to predict the impact of each variant. VEP assigns one of four IMPACT ranks (high, moderate, low and modifier) along with predicted consequences to each variant overlapping a transcript or a regulatory feature. The strength of association between the genotype at each variant and the expansion propensity phenotype was taken as the one-sided p-value of the F statistic for an ANOVA model with genotype as categorical predictor variable using R. 24 SV loci were filtered out due to not returning an association value for a final count of 9,103 SNPs, 160 STRs and 959 SVs. There were an average of 4.9 transcripts and 9.7 regulatory features per gene for a total of 398 features and 44,493 variant feature pairs. The variant-feature pair with the most severe impact and consequence was selected among variants predicted to have multiple consequences and/or impacts on protein features.

### Pangenome analysis of structural variants

The BXD pangenome for chromosome 13 was built from data of 148 strains (four strains were excluded due to poor assembly quality) using haploid assemblies of 10X reads obtained by Supernova^43^. To restrict the analysis to chromosome 13, haploid assemblies were mapped against the GRCm38/mm10.fa reference genome using wfmash65 v.0.6.0^44^. Only assemblies mapping to chromosome 13 were used to build the the pangenome with pggb^45^ v0.2.0 using the following combination of parameters: `pggb-0.2.0 -i chr13.pan+ref.fa.gz -o chr19.pan+ref -t 48 -p 98 -s 100000 -n 140 -k 229 -O 0.03 -T 20 -U –v -L -Z`.

Regions of the pangenome with depth<10x were removed using odgi^46^. Variant calling from the pangenome was done with vg^47^ (v1.35.0-59-ge5be425c6) using the following combination of parameters: `vg-e5be425 deconstruct -t 16 -P REF -e -a -H ‘#’ graph.gfa > graph.vcf`.

The variant call set was processed to remove missing data, sites where alleles are stretches of Ns, homozygous reference genotypes and variants smaller than 50bp and >10kbp before normalization and decomposition using bcftools^48^ under standard parameters. The resulting VCF file was visualized using bandage v. 0.8.1^49^.

Reference and alternate allele sequences for structural variants were extracted from the resulting variant call file using ‘bcftools query’. Each alternate sequence was then aligned to the reference using the Needleman-Wunsch global pairwise alignment implemented in the ‘pairwiseAlignment’ function from the Biostrings v2.60.1 R package. This allowed for splitting complex structural variant sequences spanning multiple kilobases into smaller individual insertions/deletions for variant effect analysis. We removed singleton variants and those less than 50bp in length.

### eQTL analysis

We generated a list of 264 expression dataset files available from GeneNetwork’s Interplanetary File System (IPFS) node (**URLs**) using the “lftp” tool. Of these, 242 datasets contained BXD strain data. A number of GeneNetwork datasets do not reflect the nomenclature change of the BXD24/BXD24_Cep sister strains. To avoid ambiguity and standardize strain names with newer datasets, BXD24 and BXD24a were relabeled as BXD24_Cep and BXD24 respectively in datasets GN267, GN373, GN385, GN410 and GN414, which contained expression values for both of these strains. Similarly, BXD24a was relabeled as BXD24 in datasets GN274, GN275, GN302, GN308, GN325, GN374, GN375, GN387 and GN702. Probe information and per-strain gene expression values were extracted into separate tables of an sqlite3 database to facilitate querying. Probes with missing genomic location information were removed. Finally, probe coordinates were converted from the mm9 to the mm10 reference using the UCSC Genome Browser liftOver tool and probes that failed remapping to the new reference were discarded.

Each GeneNetwork dataset represents a distinct processing configuration of data generated from an experimental study. Processing steps include: signal intensity normalization, strain and probe filtering, rescaling and correction of batch effects. Multiple datasets may be available for studies where both gene- and exon-level data has been collected. Further, study data may be split up into multiple datasets according to the sex of the animals or by treatment group such as diet or drug exposure. To avoid overcounting, we selected a single representative dataset by using a heuristic approach to make the selection based on strength of signal and processing conditions. Exon level data was preferred to gene level data due to increased probe density. More recently reprocessed datasets were preferred to older ones. Data from control groups was preferred to data from experimentally treated groups. Combined male and female data was preferred to sex-specific data. Datasets with more strains were preferred to datasets with fewer strains. A summary of selected and available datasets for each study is available in **Supplementary Table 4**.

We then queried expression values for all probes falling within the 83.8-93.4 Mb region on chromosome 13 in each dataset. GN227 lacked probe data in this region and was excluded. Probe mapping information was either taken directly from the GeneNetwork dataset or queried from Ensembl’s BioMart data mining tool release 102 using the biomaRt^50^ R package. Unmapped probes were removed from analysis. We then checked whether probe coordinates were contained within the start and stop positions of each probe’s corresponding gene and removed those that did not. For each Affymetrix ProbeSet representing a collection of probes, we used the UCSC^51^ BLAT tool (https://genome.ucsc.edu/goldenpath/help/blatSpec.html) to find the matching genomic location of individual probe sequences. We discarded probe sets where any contained probe did not match within the coordinates of its assigned gene. We then used probe coordinates to calculate the number of segregating variants that each probe overlapped using the ‘bedtools intersect’ command available from the BEDTools^52^ package. Additionally for each probe, we calculated the number of variants at which each strain differed from the mm10 reference, which represents the number of mismatches an array probe would be expected to have when hybridizing with a DNA library sample from a given strain. We then performed eQTL mapping on chromosome 13 using the same set of LD-independent loci and kinship matrix. The covariate vector from the QTL mapping was supplemented with the number of expected hybridization mismatches for each probe/strain combination to account for expected differences in hybridization efficiency. The number of strains per dataset ranged from 11 to 79. For comparison, we remapped the mutation propensity phenotype using only strains available in each of the gene expression datasets. Mono-allelic markers conditioned on the subset of strains available in each expression dataset were removed.

Notably, it is common for multiple microarray probes (probesets) to target the same gene, especially for exon based microarrays. We observed high variability for gene expression measurements between probes targeting the same gene in a given dataset. To limit the rate of false eQTL signal discovery, we applied the Benjamini Hochberg (https://www.jstor.org/stable/2346101) multiple hypothesis testing correction to the vector of peak LOD values for each gene-dataset pair. We selected a representative probe for each gene having the highest adjusted peak LOD value within the expansion phenotype QTL region on chr13 for gene-level analysis. For visualization of eQTL traces, LOD values at each marker were scaled by the ratio of the peak adjusted LOD to peak LOD for each gene

### Genomic data for classic mouse strains

Read alignment bam files for the common laboratory mouse strains: 129S1/SvImJ, NZO/HILtJ, NOD/ShiltJ, CAST/EiJ, PWK/PhJ, A/J and WSB/EiJ were downloaded from the Mouse Genomes Project ftp server hosted at ftp://ftp-mouse.sanger.ac.uk/current_bams. Variant call files for these strains were similarly queried from ftp://ftp-mouse.sanger.ac.uk/current_snps.

### Tissue-specific expression of DNA repair genes

Tissue-specific expression of *Msh3* and other DNA repair genes (**Supplementary Table 5**) was obtained from the Bgee database^32^, accessed on February 26, 2022.

## Data availability

Whole genome sequencing data and genotype calls for the 152 RI strains from BXD will be released upon publication of Ashbrook et al^17^.

## Code availability

Workflow and analysis scripts are available at: https://github.com/mikmaksi/BXD_str_expand_prop_pub

## Acknowledgements

This work was supported by NIH/NIDA award 1U01DA051234 (A.A.P., M.G., and J.S.).

## URLs

BXD Genotype Markers: http://gn1.genenetwork.org/webqtl/main.py?FormID=sharinginfo&GN_AccessionId=600. GeneNetwork Datasets: http://ipfs.genenetwork.org/ipfs/QmakcPHuxKouUvuNZ5Gna1pyXSAPB5fFSkqFt5pDydd9A4/ *Msh3* expression patterns: https://bgee.org/?page=gene&gene_id=ENSMUSG00000014850

## Author contributions

M. M. performed and led data analysis and wrote the paper. D. G. A. and the BXD Sequencing Consortium generated WGS for BXD strains and gave feedback on analyses. D. G. A. additionally helped with analysis and access of GeneNetwork expression datasets. F. V. and V. C. performed pangenome analysis of structural variants. N. Mo. helped with analysis of STR VCF files. N. Ma. helped with analysis of the LTR element in classic mouse strains. A. A. P. and M. G. obtained funding for the project, supervised analyses, and wrote the paper.

