## Supplementary Figures for "A novel quantitative trait locus implicates *Msh3* in the propensity for genome-wide short tandem repeat expansions in mice"

Supplementary Fig. 1

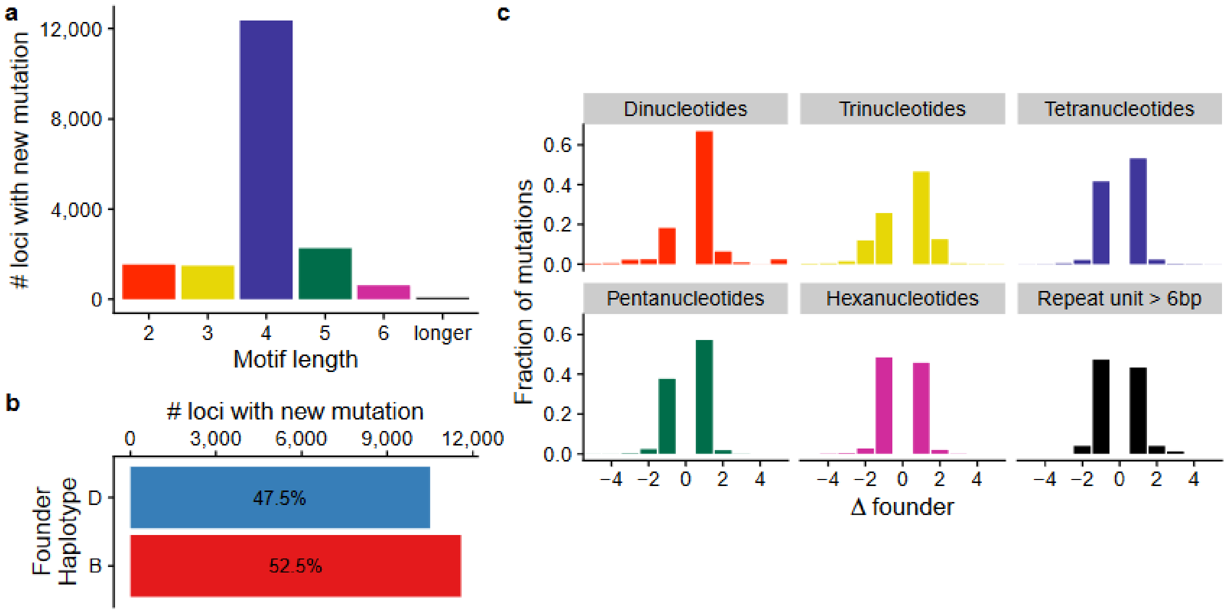


**Summary of new STR mutations in BXD. a. Distribution of repeat unit lengths among new STR mutations.** The number of new mutations at STRs with each repeat unit length (bp) is shown. **b. Distribution of the founder haplotype for new mutations.** Bars show the number of new STR mutations occurring on "B" (red) vs. "D" (blue) founder haplotypes. **c. Distribution of mutation sizes for each repeat unit length.** The x-axis shows mutation sizes in terms of the difference in number of repeat units (RU) from the founder allele. Positive sizes indicate expansions and negative sizes indicate contractions.

Supplementary Fig. 2

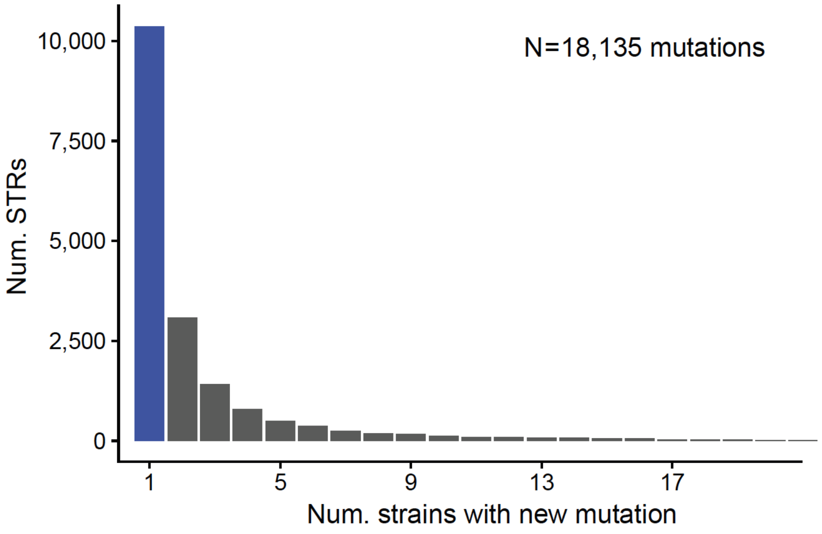


**Distribution of the number of strains carrying the new allele at each of the STRs for which at least one new mutation was identified.** Data is shown for N=18,135 unique STRs. Singleton mutations, seen only in a single strain, are shown in blue.

### Supplementary Fig. 3

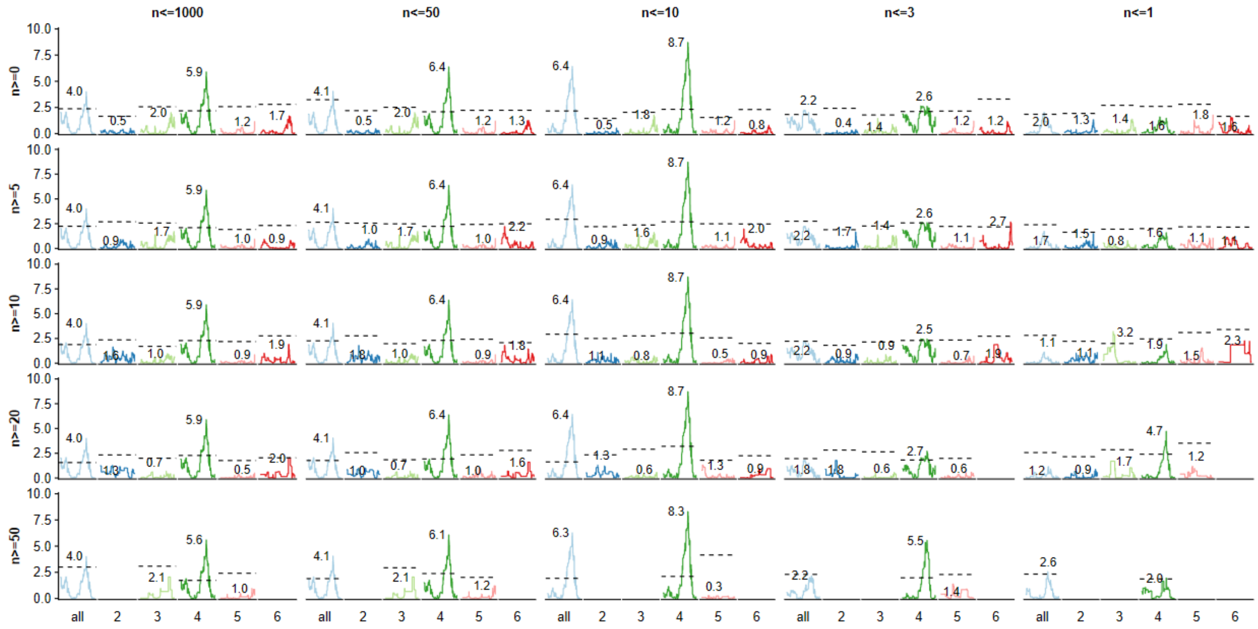


**Evaluating robustness of the chromosome 13 association signal for expansion propensity.** In each panel, the x-axis denotes the repeat class (from left to right: all STRs, and including only STRs with a repeat unit length of 2-6bp). Within each class in each panel, the x-axis denotes genomic location on chromosome 13 and the y-axis denotes logarithm of the odds (LOD). The max LOD is annotated for each class. Each row denotes a different threshold for the minimum number of new STR mutations for a strain to be included in the analysis (strain filtering). Each column denotes a different threshold for filtering the maximum number of strains a particular new STR mutation could be observed in (frequency filtering). Dashed horizontal lines represent permutation thresholds for genome-wide significance in each class. Overall, strain filtering has little effect whereas frequency filtering indicates the association signal is restricted to relatively new mutations. In all cases, tetranucleotides, the largest STR class in our dataset, show the strongest signal.

Supplementary Fig. 4

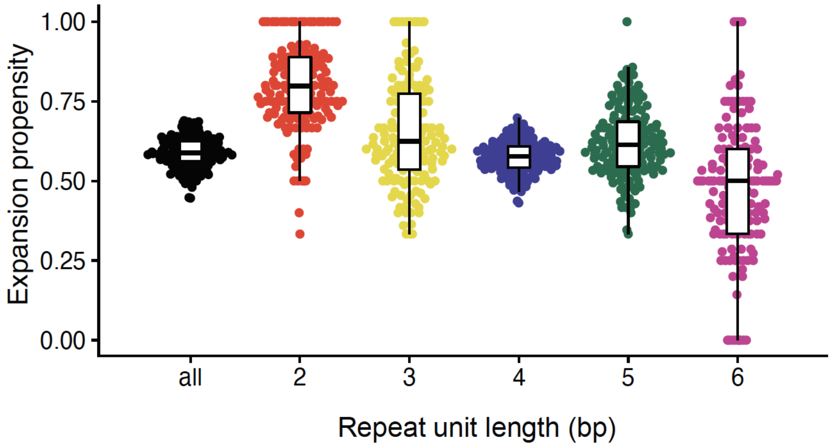


**Distribution of expansion propensity for each strain for different repeat classes.** Expansion propensity was computed separately considering only STRs with repeat units of a specified length (black=all STRs; red=dinucleotides; gold=trinucleotides; blue=tetranucleotides; green=pentanucleotides; purple=hexanucleotides).

Supplementary Fig. 5

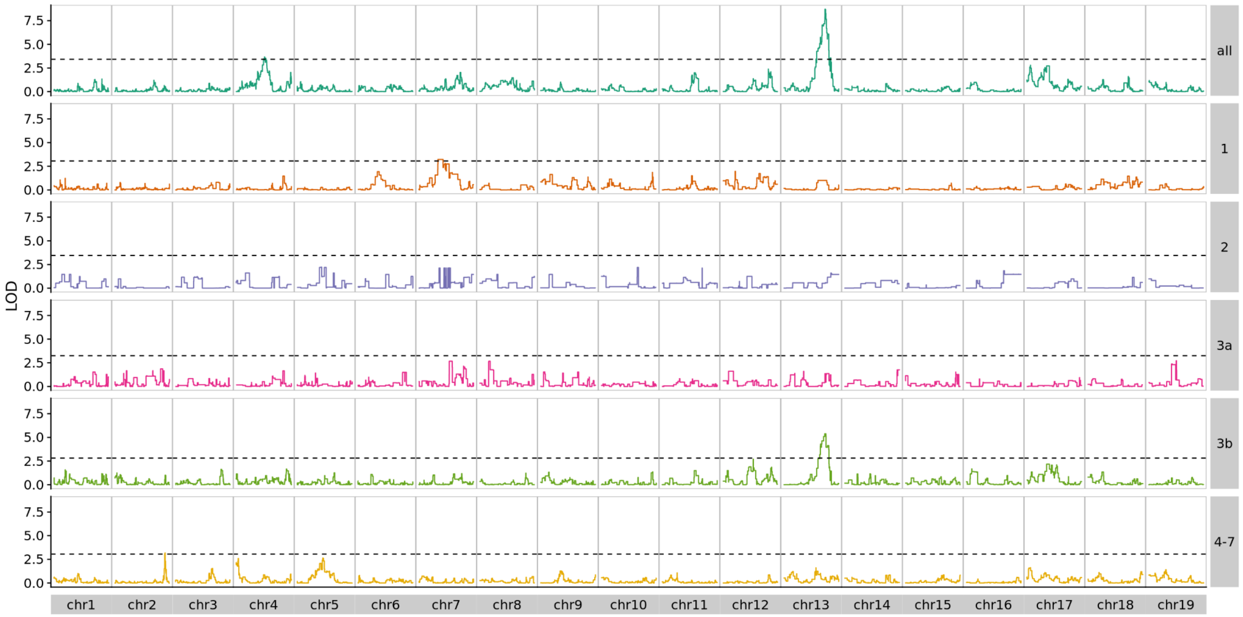


**Expansion propensity QTL mapping in each BXD epoch.** We repeated QTL mapping separately using only strains in each epoch. Each row represents a different epoch. In each row, the x-axis denotes genomic location and the y-axis denotes LOD score. Permutation based thresholds are shown as dashed horizontal lines.

Supplementary Fig. 6

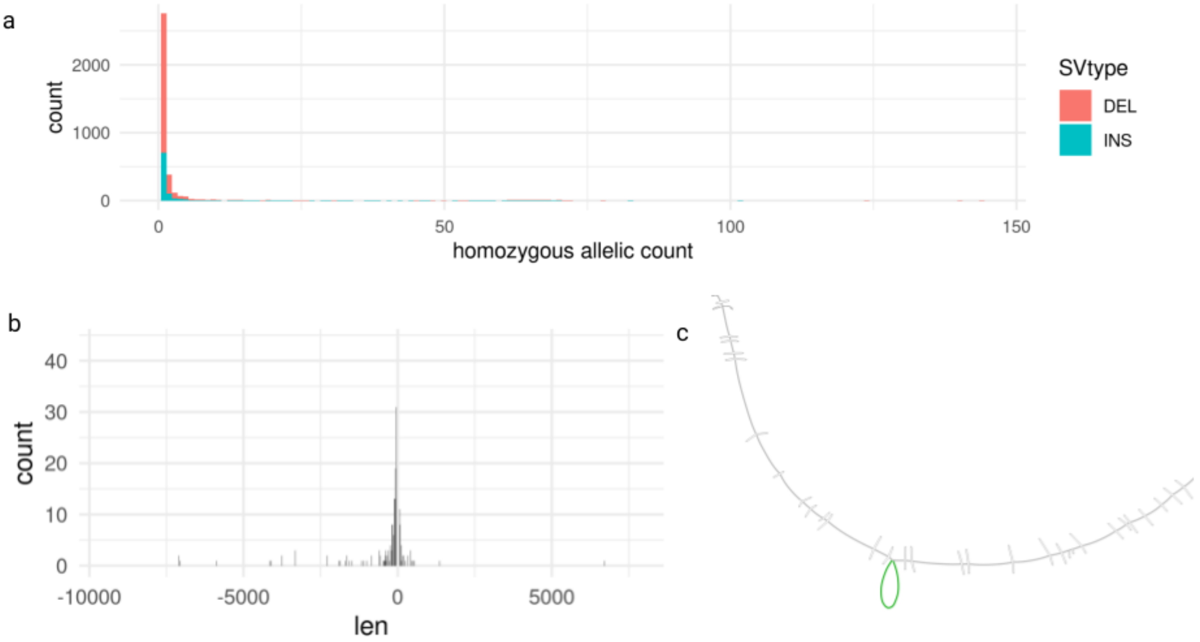


**Features of the SVs discovered from pangenome analysis of chromosome 13. a.** Allele frequency spectrum of the 3,698 SVs with length >50bp and <10kbp in the region of interest (chromosome 13 region 87,348,000-97,348,498). **b**. Distribution of the length of insertions and deletions. **c.** Bandage[1] representation of the candidate region on chromosome 13 (mm10, chr13:92,345,000-92,351,498) containing the 387bp insertion found in the 66 mice with C57BL/6J background for that region.

Supplementary Fig. 7

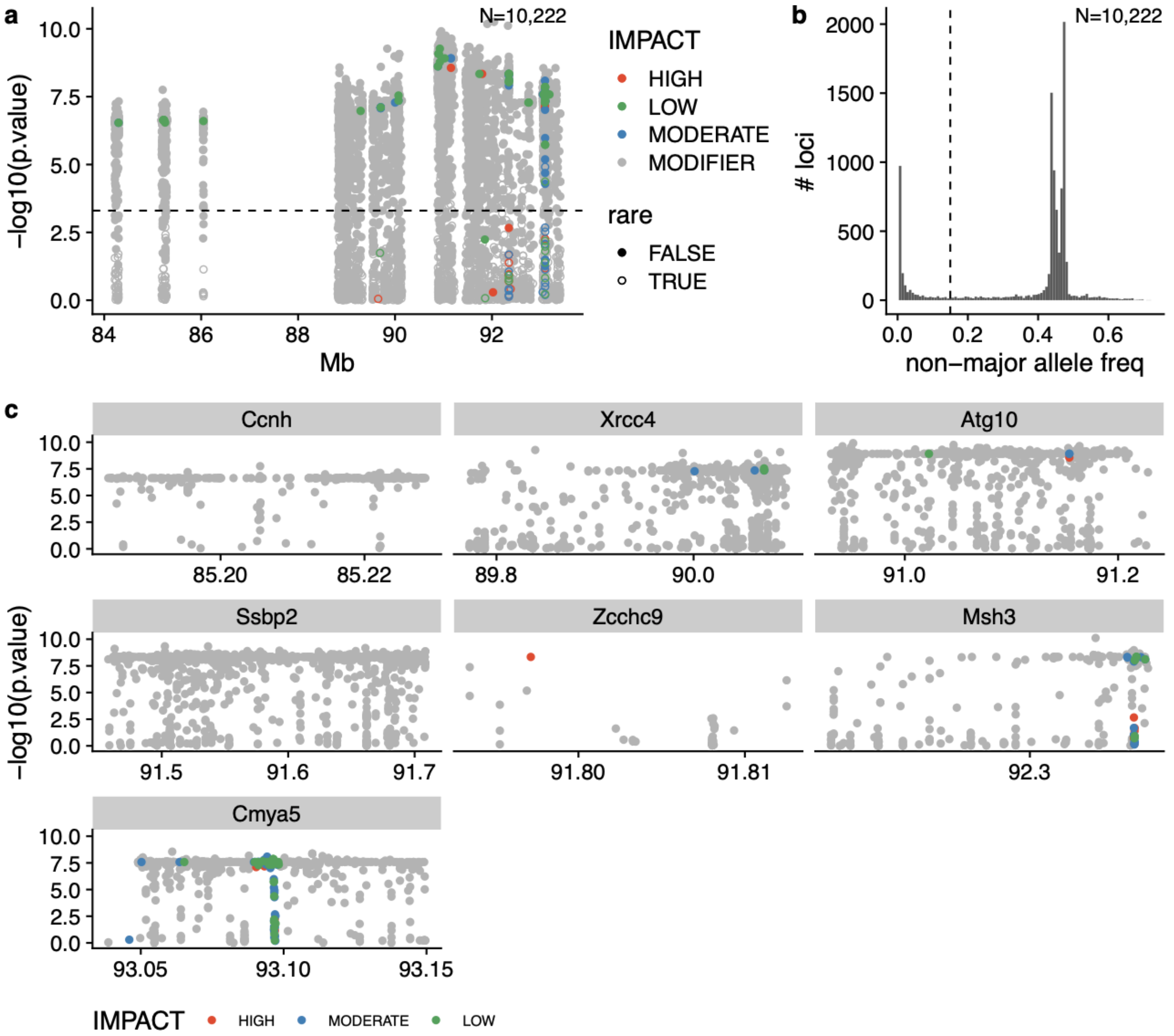


**Annotation and selection of impactful variants within genes in the chromosome 13 QTL for expansion propensity.** For plots in **a** and **c**, the x-axis gives the genomic coordinate and the y-axis gives the association (-log10 p-value). Each dot represents a variant, and variants are colored by their impact predicted by VEP (red=high, blue=moderate, green=low; gray=modifier). **a. VEP-annotated variants across the entire QTL region.** Most annotated variants are located in intronic regions and have a predicted “modifier” impact. Weakly associated variants were removed from further analysis using a threshold of 3.3 on the association statistic (dashed horizontal line) as suggested on the GeneNetwork website (http://gn1.genenetwork.org/glossary.html). **b. Distribution of non-major allele frequencies.** Rare variants with an artificially strong association statistic due to overleveraging of outliers were removed using a threshold (dashed vertical line) on non-major allele frequency. **c. Detailed view of VEP-annotated variants.** Views are shown for genes known to be involved in DNA repair (*Ccnh*, *Xrcc4, Atg10, Ssbp2, Msh3*) or genes for which high-impact variants were detected (*Cmya5, Zcchc9*).

### Supplementary Fig. 8

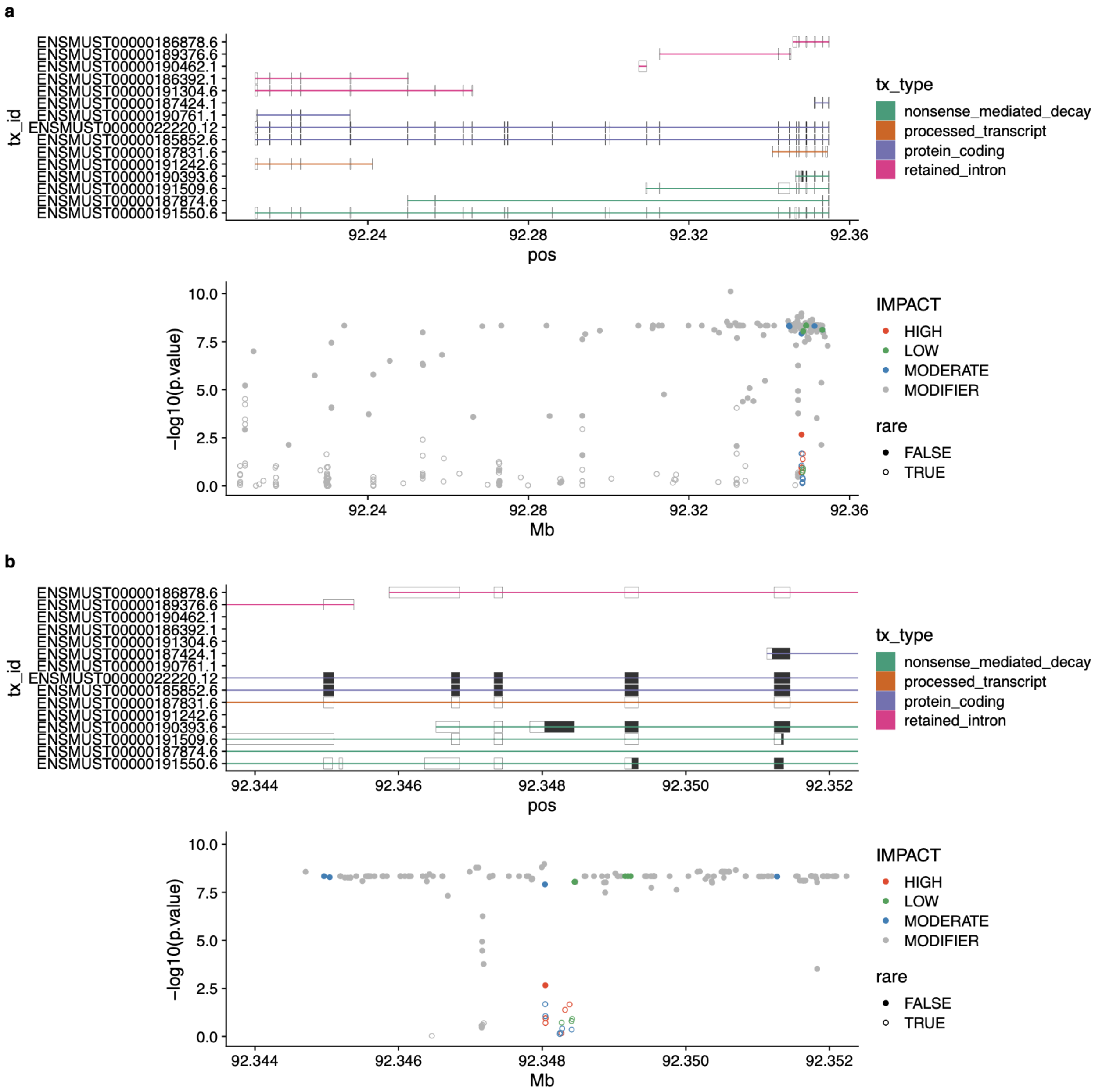


**Detailed view of annotated variants within *Msh3*.** In **a.** and **b.**, top panels show transcript annotations, colored by transcript type. In bottom panels, the x-axis gives the genomic coordinate and the y-axis gives the -log10 p-value of each variant for association with expansion propensity. Variants are colored by VEP-predicted impact. Filled dots represent common and empty circles represent rare variants based on the threshold identified in **Supplementary Fig. 7b.** Plots are the same as those in **Fig. 3**, but include additional transcript annotations and rare variants. **a.** Shows the entire length of *Msh3*, whereas **b.** zooms in on the variant-dense 5' region. High-impact rare variants overlap a 387bp LTR insertion in the “B” haplotype and likely represent variant calling artifacts.

Supplementary Fig. 9

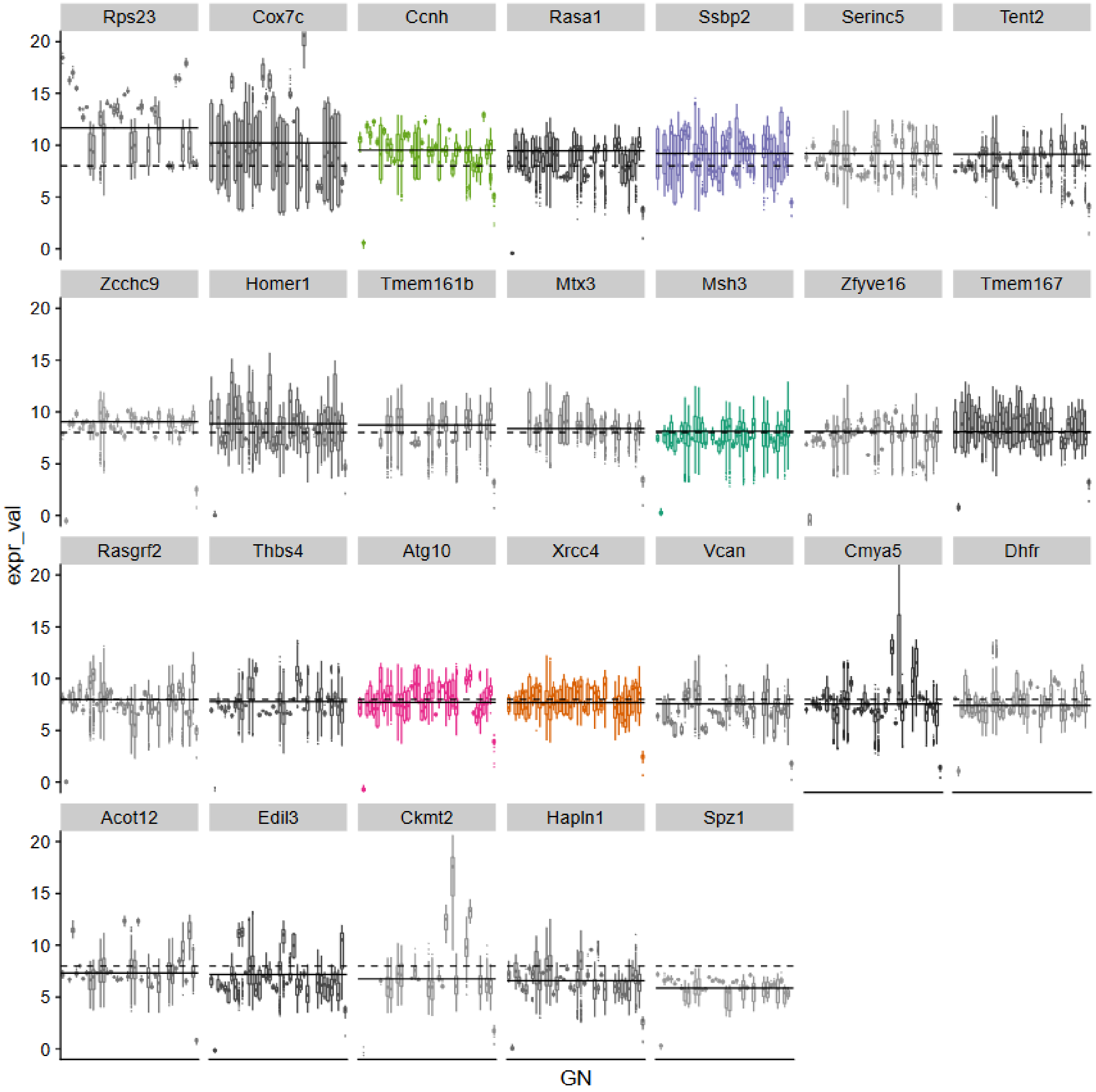


**Overall gene expression levels for genes with the QTL region.** Boxplots show distributions of normalized gene expression levels for each of the protein coding genes within the QTL confidence interval. Each gene is shown in a separate panel. Distributions are ordered by GeneNetwork dataset id (x-axis) and panels are ordered by the median gene expression level across all datasets (solid horizontal line). GeneNetwork datasets are normalized using a "2z+8" method[2]. The expected average value of 8 is shown as a dotted horizontal line.

Supplementary Fig. 10

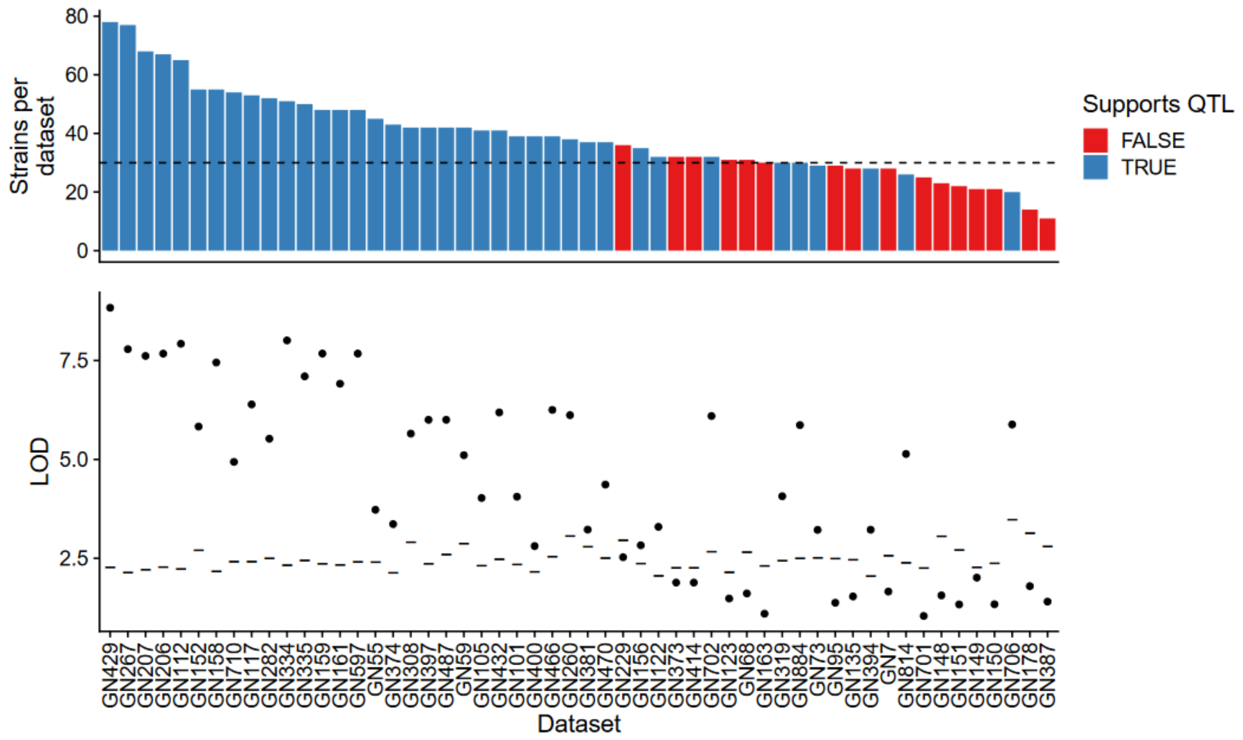


**Summary of expansion propensity QTL signal detection using strains available in each gene expression dataset.** The top panel shows the number of strains included in each expression dataset. Datasets are sorted in decreasing order by the number of strains per dataset. The dashed line indicates the minimum strain-per-dataset cutoff of 30 strains. We performed QTL analysis for expansion propensity using the subset of strains available in each expression dataset. The bottom panel shows peak LOD (black points) for each dataset. Gray dashes show the permutation-based significance threshold computed separately for each dataset. Blue bars in the top panel indicate the subset of strains available in that expression dataset was sufficient to reproduce the QTL for expansion propensity.

Supplementary Fig. 11

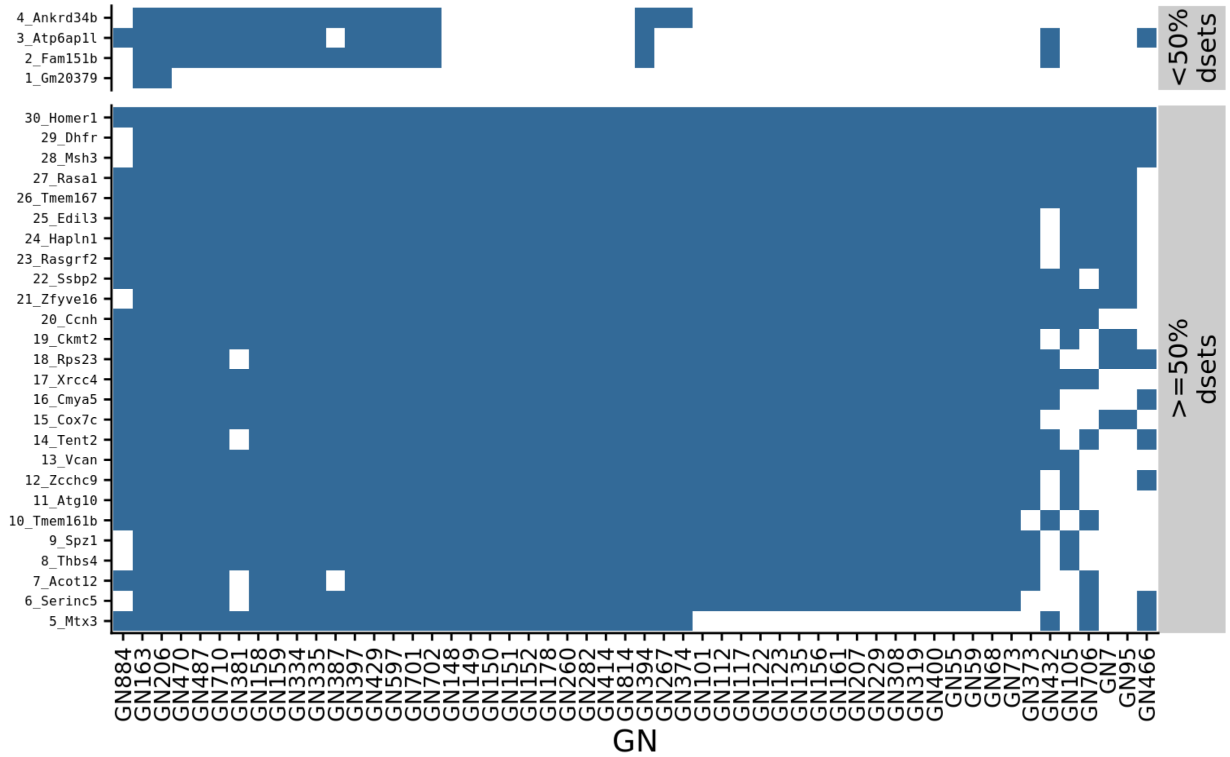


**Availability of gene expression data for genes within the expansion propensity QTL.** Of the 30 protein-coding genes shown, 26 (bottom panel) had gene expression values in at least 50% of the representative microarray datasets (x-axis) selected from GeneNetwork and 4 did not (top panel).

Supplementary Fig. 12

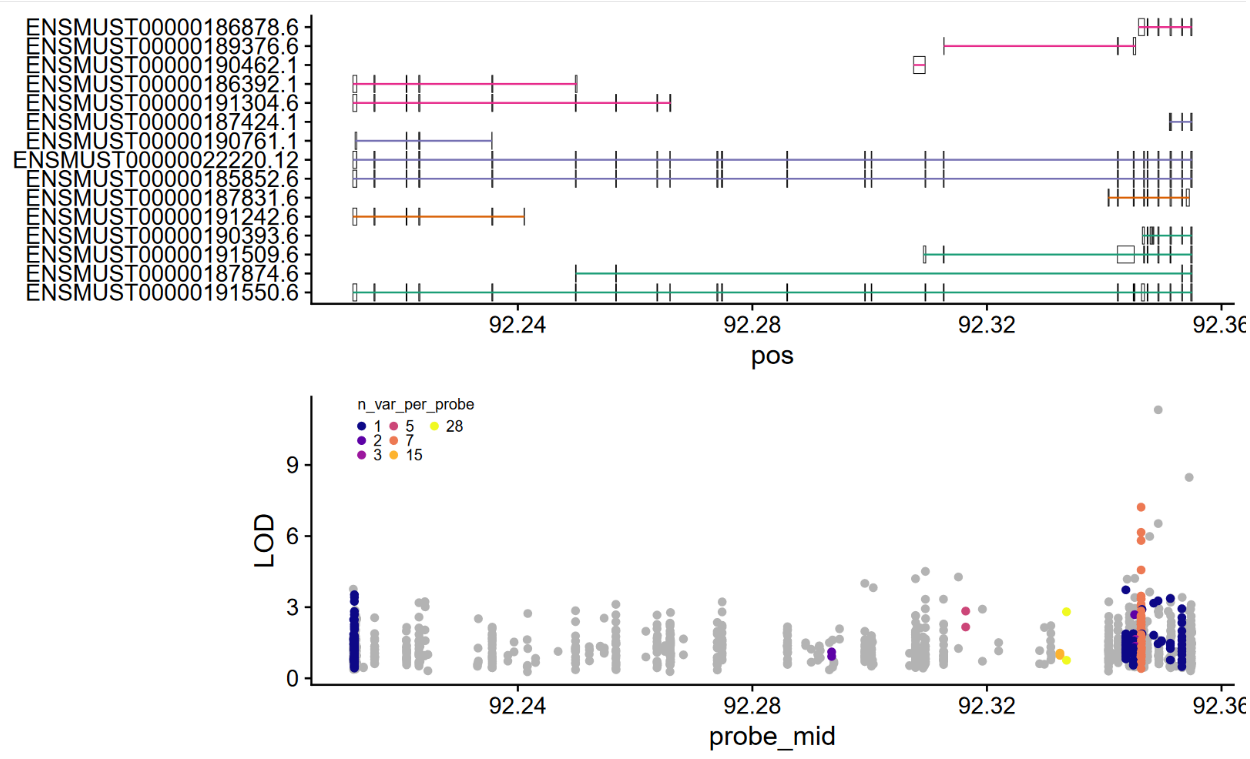


**Probe-level analysis of eQTL signals at *Msh3*.** The top panel annotates *Msh3* transcripts. In the bottom panel, each dot represents a single microarray probe. The x-axis gives the position of each probe. The y-axis gives the maximum LOD score across all datasets for each probe. Probes are colored by the number of segregating SNPs overlapping the probe coordinates. Probes not overlapping SNPs are shown in gray. Probes near the 5' end of *Msh3* showed the strongest eQTL signals. However the majority of those overlap SNPs, which could lead to biased expression measurements and were filtered from gene expression analysis

Supplementary Fig. 13

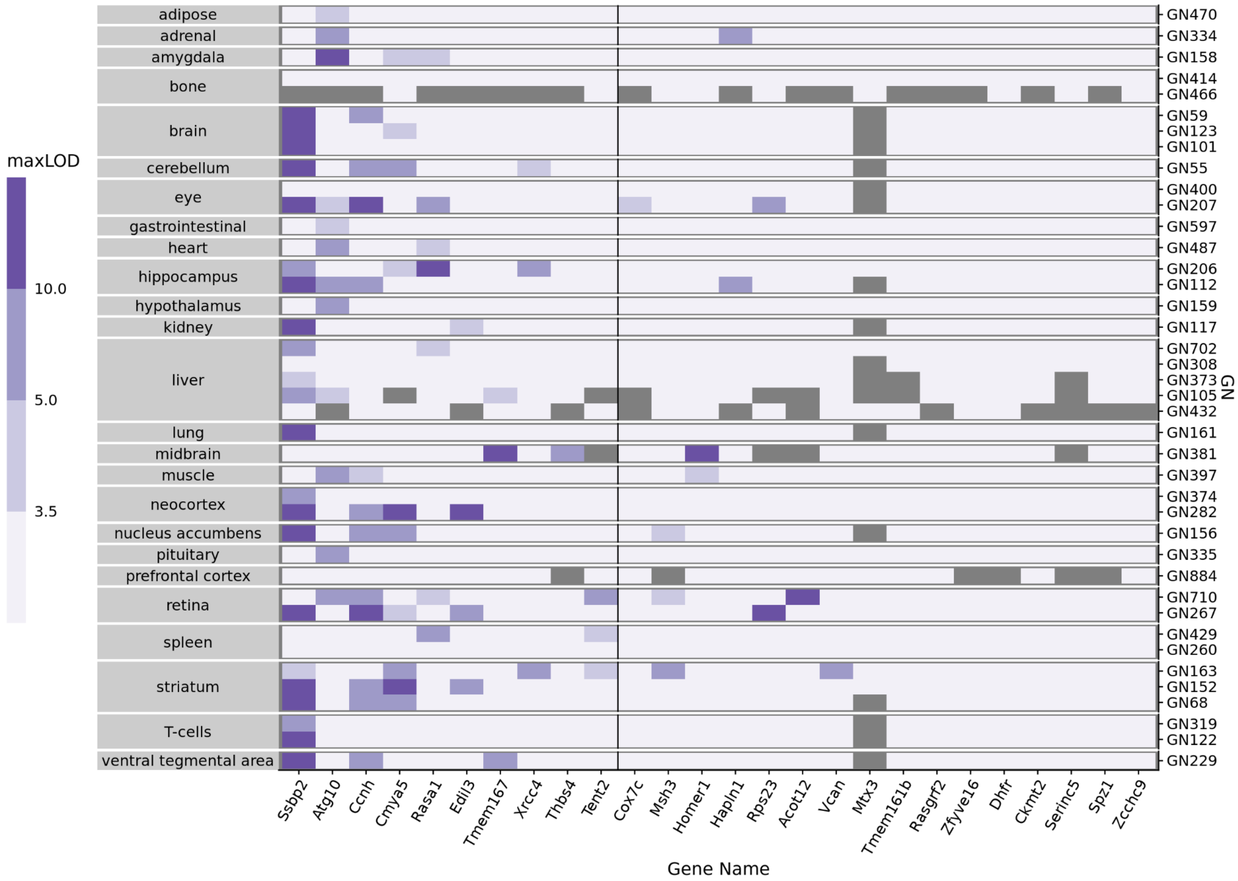


**Summary of gene eQTL signals for genes contained within the QTL peak 1.5-LOD support interval for the expansion propensity phenotype.** eQTL mapping was performed for each probe corresponding to a gene within the region of interest compiled across all GeneNetwork datasets. The maximum LOD value is shown for each gene (columns) in each dataset (rows, grouped by tissue). Genes are ordered from left-to-right according to the number of datasets in which the peak LOD eQTL value exceeded the permutation based threshold in that dataset. The vertical black line denotes the top 10 genes. While a single dataset is available for most tissues (primary y-axis), multiple independent datasets are available for others. GeneNetwork dataset ids are shown on the right y-axis.

Supplementary Fig. 14

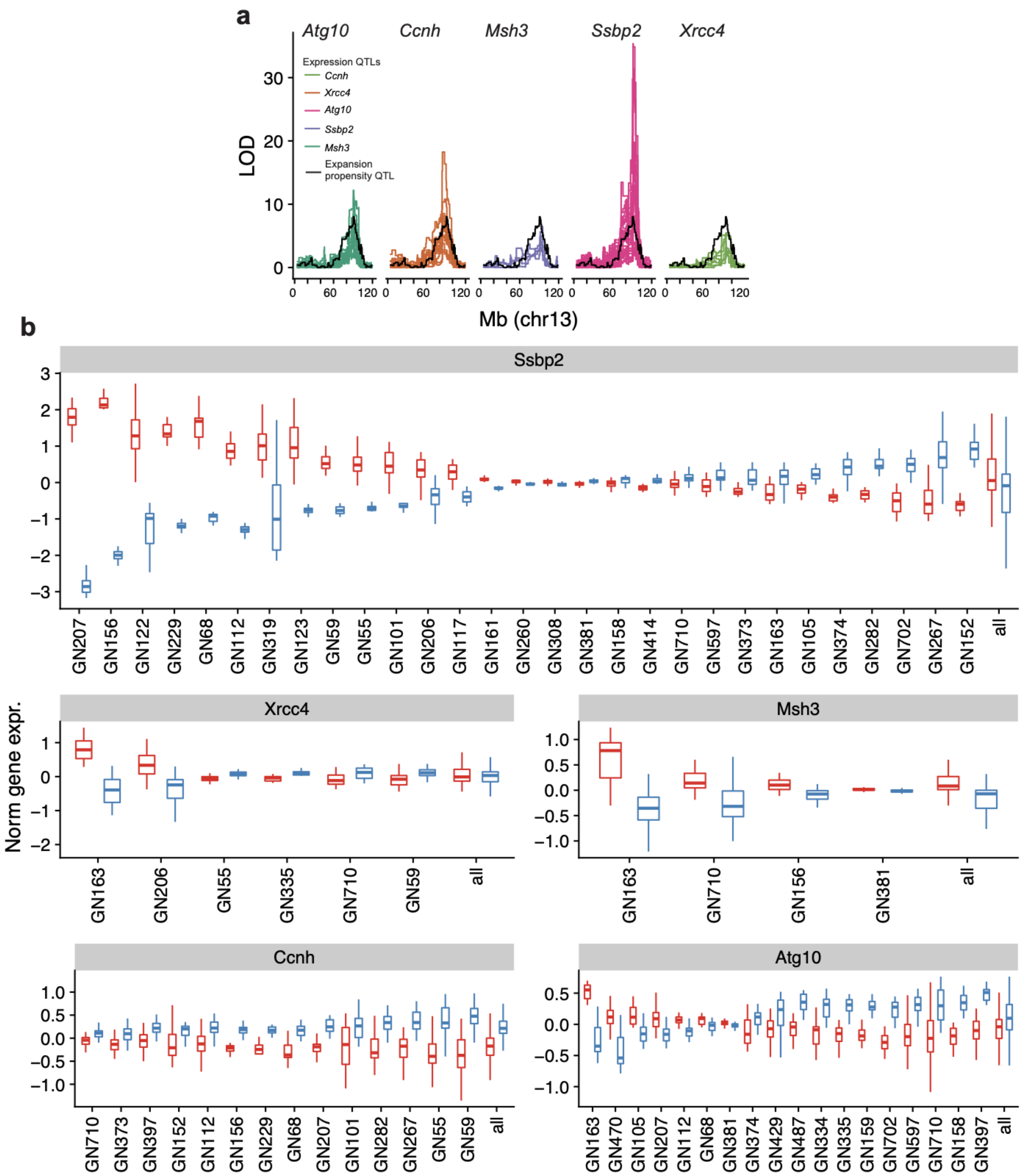


**eQTL signals for genes within the expansion propensity QTL. a. Co-localization of expansion propensity and eQTL signals.** LOD scores for expansion propensity are shown in black. Colored traces denote eQTL LOD scores. A separate line is shown for each expression dataset. **b. Distribution of gene expression for strains with "B" vs. "D" haplotypes.** Panels show gene expression at DNA-repair related genes for strains assigned the "B" (red) vs. "D" (blue) haplotypes at the chr13 expansion propensity locus. Each column denotes a different GeneNetwork expression dataset (**Supplementary Table 2).** Datasets are ranked by the difference in expression between strain groups. Only datasets where a significant eQTL was identified are shown. The far right column shows data aggregated across expression datasets.

Supplementary Fig. 15

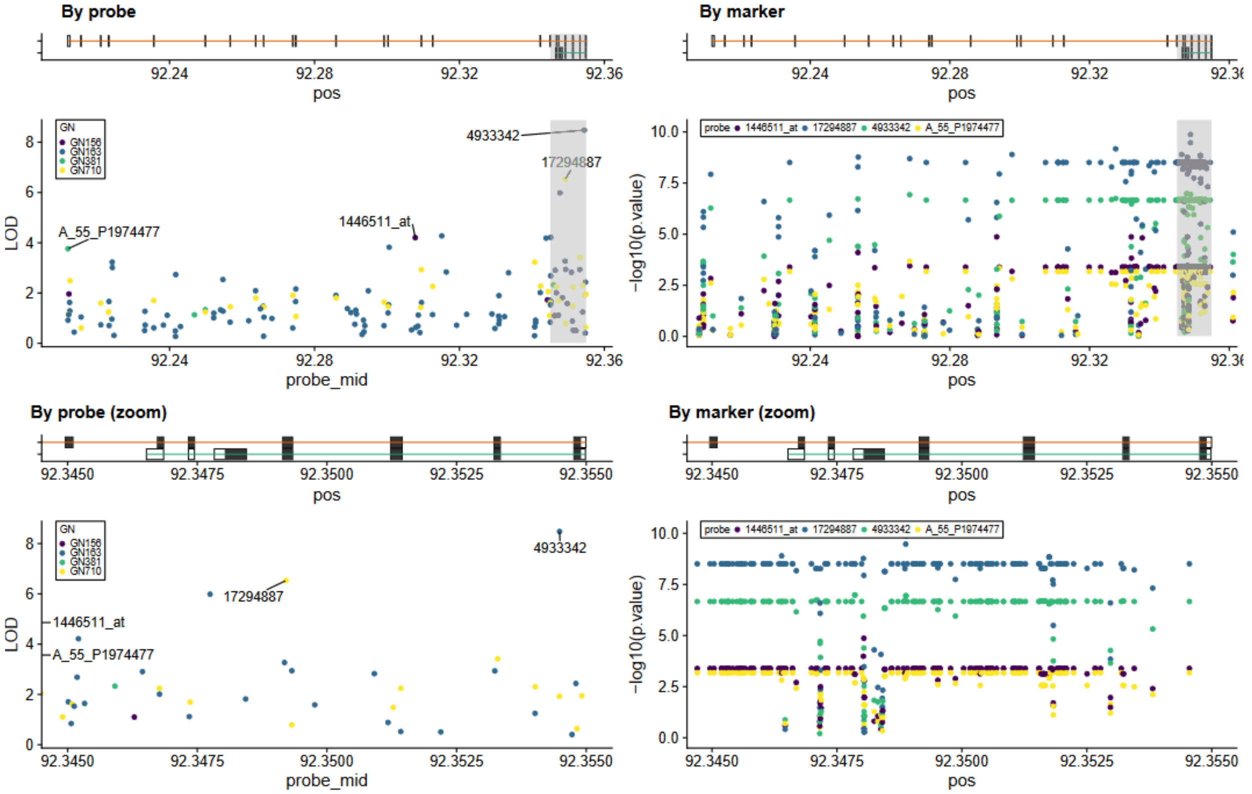


**Detailed analysis of eQTL signals at *Msh3*.** Left panels show the location of each microarray probe (x-axis) and the maximum LOD score across all variants for association with that probe. Colors represent different GN datasets. Right panels show the location of each variant (x-axis) and the best -log10 p-value across all *Msh3* probes. Colors denote different microarray probes. Bottom panels show zoomed-in views denoted by the gray rectangles in top panels, which contain both the probes and variants with the strongest eQTL signals near the 5' end of *Msh3*.

Supplementary Fig. 16

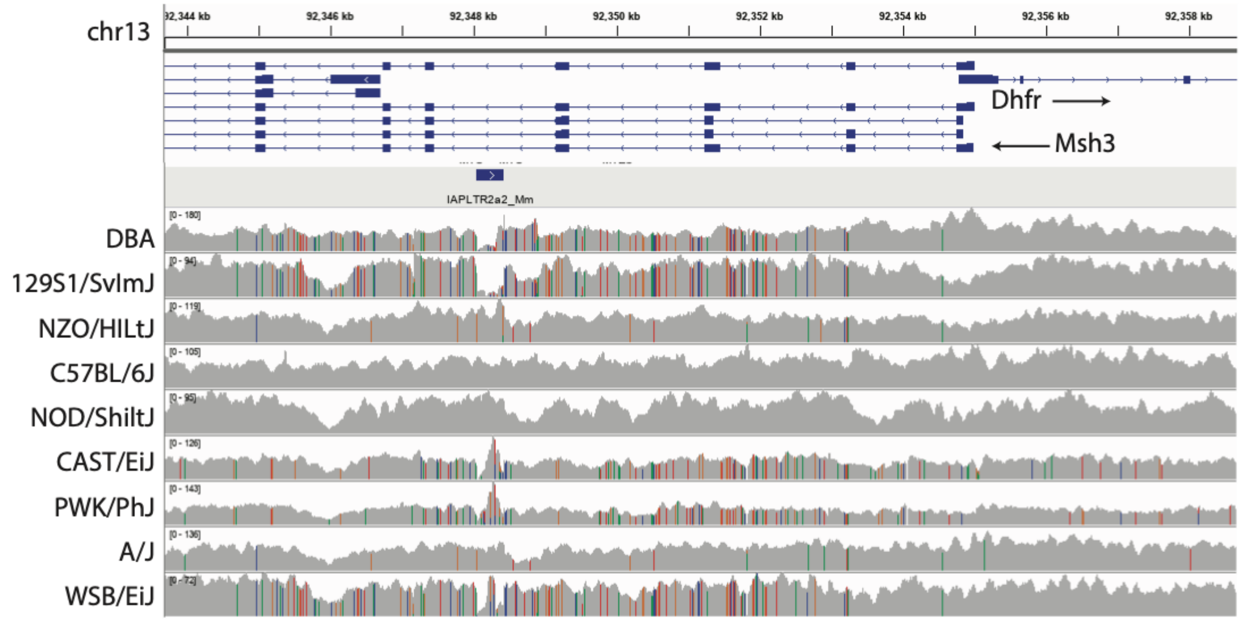


**Visualization of next-generation sequencing data for classic mouse strains at the 5' end of *Msh3*.** Top tracks show gene annotations. The middle track denotes the location of the IAP LTR element described in the main text. Bottom tracks show sequencing coverage in classic mouse strains. Colored bars indicate sequence variants compared to the mm10 reference genome, which is based on C57BL6/J. Strains 129S1/SvImJ and WSB have similar haplotypes in this region to DBA ("D"), whereas NOD is similar to C57BL6/J. Coverage profiles suggest strains DBA, 129S1/SvImJ, CAST/EiJ, PWK/PhJ, and WSB/EiJ do not have the IAP LTR insertion present in the reference genome. The visualization was created using the Integrative Genomics Viewer (IGV).
